# Locityper: targeted genotyping of complex polymorphic genes

**DOI:** 10.1101/2024.05.03.592358

**Authors:** Timofey Prodanov, Elizabeth G. Plender, Guiscard Seebohm, Sven G. Meuth, Evan E. Eichler, Tobias Marschall

## Abstract

The human genome contains numerous structurally-variable polymorphic loci, including several hundred disease-associated genes, almost inaccessible for accurate variant calling. Here we present Locityper, a tool capable of genotyping such challenging genes using short and long-read whole genome sequencing. For each target, Locityper recruits and aligns reads to locus haplotypes, for instance extracted from a pangenome, and finds the likeliest haplotype pair by optimizing read alignment, insert size and read depth profiles. Locityper accurately genotypes up to 194 of 256 challenging medically relevant loci (95% haplotypes at QV33), an 8.8-fold gain compared to 22 genes achieved with standard variant calling pipelines. Furthermore, Locityper provides access to hyperpolymorphic HLA genes and other gene families, including KIR, MUC and FCGR. With its low running time of 1h10m per sample at 8 threads, Locityper is scalable to biobank-sized cohorts, enabling association studies for previously intractable disease-relevant genes.

## Introduction

Single-nucleotide variants (SNVs) are the most abundant class of genetic variants segregating in the human population and are at the same time easy to access using microarray or short-read sequencing platforms. Unsurprisingly, virtually all genome-wide association (GWAS) studies seeking to map genotypes to phenotypes have therefore been focusing on SNVs. In contrast, structural variants (SVs), which are 50bp in size or longer, are much more challenging to characterize and more than half of all SVs per sample are missed by short-read based variant discovery ^1–3^, despite their biomedical relevance ^4,5^. This difficulty to analyze SVs from short reads is largely driven by their common formation through homology-associated mechanisms, leading to repetitive and complex sequence contexts ^2^. Almost 750 genes contain “dark” protein-coding exons, where read mapping and variant calling cannot be adequately performed ^6^ and around 400 medically relevant genes are almost inaccessible due to their repetitive nature and high polymorphic complexity ^7^. Of them, 273 genes are widely used for variant calling and assembly benchmarking ^8,9^. Long read technologies are needed to address this problem ^10–12^ and recent long-read based genome assembly strategies indeed lead to haplotype-resolved genome assemblies of diploid samples that routinely resolve many previously intractable complex genetic loci ^13,14^. Nevertheless, long read sequencing of large cohorts remains prohibitively expensive, signifying the need for accurate short read based genotyping.

In the meantime, high quality assemblies are available for hundreds of human haplotypes and give rise to a pangenome reference ^2,8,15^. The genetic variation encoded therein can serve as a basis for genotyping workflows by mapping reads to a pangenome graph ^16,17^ or through *k*-mer based genome inference ^18^. While genome inference with Pangenie ^18^ has expanded the set of accessible structural variants considerably ^8^, it exhibits limitations at complex loci with few unique *k*-mers. As an alternative strategy, methods for targeted genotyping of genes of special interest, such as the HLA and KIR gene families, have been developed ^19–21^. Even though these methods can achieve high accuracy, they typically rely on specific gene structure and cannot be easily scaled to include more targets.

Here, we propose a new tool, called Locityper, to leverage genome assemblies in a pangenome reference or custom collections of locus alleles for fast targeted genotyping of complex loci. Locityper can efficiently process both short and long read data, and it integrates a range of different signals based on read depth, alignment identity, and paired-end distance in a statistical model to infer genotype likelihoods. This provides an opportunity to genotype and analyze a diverse set of previously understudied genes for already available large sequencing datasets such as the 1000 Genomes Project cohort and large biobanks like the All-of-Us ^22^ program and the UK Biobank ^23^, where disease association studies can be performed.

## Results

### Overview of the method

Locityper is a targeted genotyping tool designed for structurally-variable polymorphic loci. For every target region, Locityper finds a pair of haplotypes (locus genotype) that explain input whole genome sequencing (WGS) dataset in a most probable way. Locus genotyping depends solely on the reference panel of haplotypes, which can be automatically extracted from a variant call set representing a pangenome (VCF format), or provided as an input set of sequences (FASTA format). Before genotyping, Locityper efficiently preprocesses the WGS dataset and probabilistically describes read depth, insert size, and sequencing error profiles. Next, Locityper uses haplotype minimizers to quickly recruit reads to all target loci simultaneously.

At each locus, Locityper estimates a likelihood for every possible locus genotype by distributing recruited reads across possible alignment locations at the corresponding haplotypes (Figure 1). The likelihood function is defined in such a way to prioritize read assignments with smaller number of sequencing errors; optimal insert sizes across the read pairs; and stable read depth without excessive dips or rises. We show that finding a maximum likelihood read assignment can be formulated as an integer linear programming (ILP) problem (Methods), for which Locityper employs existing ILP solvers and stochastic optimization. Finally, Locityper identifies genotypes with the highest joint likelihood, calculates its quality score, and outputs the most probable read alignments to the two corresponding haplotypes.

**Figure 1.**
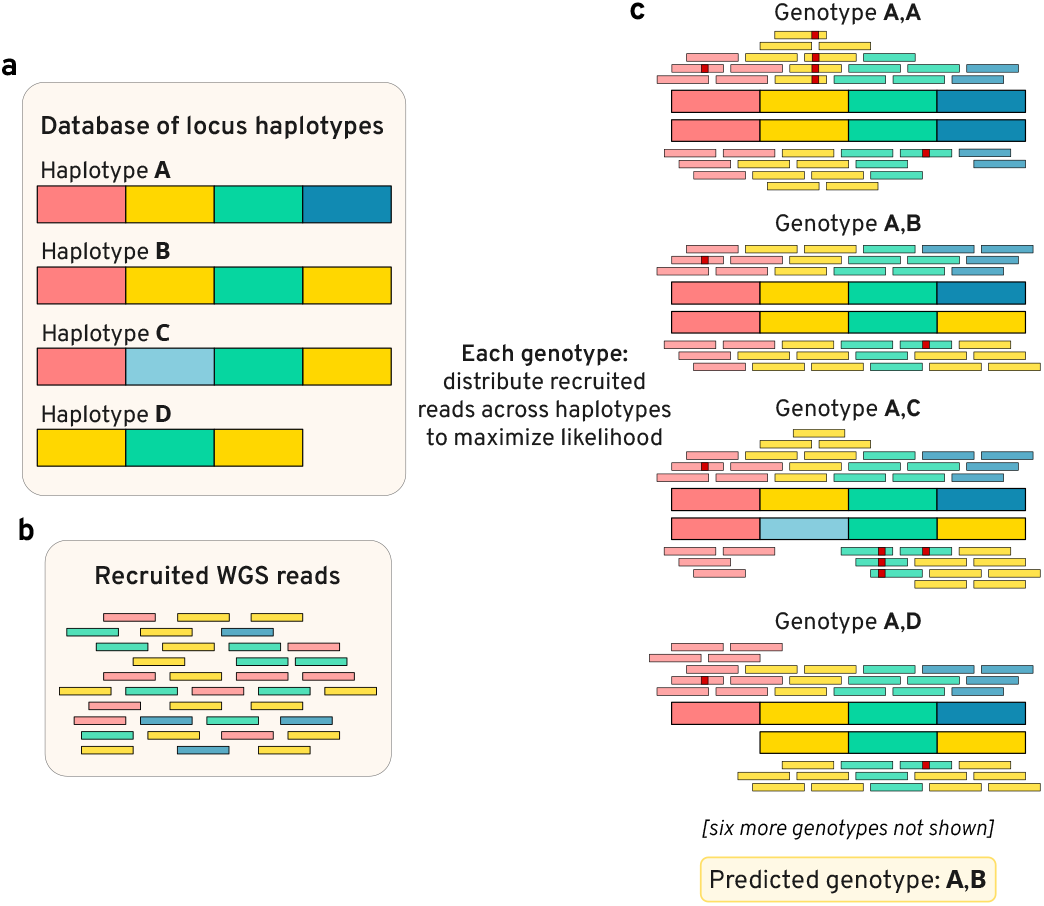
Illustration of the locus genotyping approach. **a**, Database of four locus haplotypes *(A*..*D)*. **b**, Whole genome sequencing (WGS) reads, recruited to any of the haplotypes. For illustrative purposes, haplotypes and reads are colored by homologous blocks (information, unavailable to Locityper). **c**, Optimal assignments of reads to various genotypes, where small red squares show read alignment mismatches or indels. Genotype *A*,*B* has the highest joint likelihood due to a small number of alignment errors and no lack or excess of read depth.

Locityper performs well on various sequencing technologies, including short-read Illumina sequencing, high-accuracy PacBio HiFi long reads, and error-prone PacBio CLR and Oxford Nanopore long reads. It is able to efficiently process both mapped and unmapped input reads. Across all steps, Locityper efficiently utilizes multiple computing cores to perform the analysis as fast as possible.

### Locityper accurately genotypes a wide range of challenging loci

In order to evaluate Locityper’s targeted genotyping accuracy, we utilized a list of 273 challenging medically relevant (CMR) genes, collected by Wagner et al. ^7^ For each target locus we used a reference panel of up to 90 haplotypes extracted from the recent phased whole genome assemblies ^8^ released by the Human Pangenome Reference Consortium (HPRC) ^15^. After removing genes that overlap pangenomic variants longer than 300 kb and merging nearby genes (see Methods), we retained 256 loci covering 13.9 Mb and fully encompassing 265 CMR genes and 23 other protein coding genes (see Supp. Table 1). Then, we used Locityper to genotype 40 Illumina, 40 simulated short-reads, 20 PacBio HiFi and 20 Oxford Nanopore (ONT) HPRC WGS datasets. Each of the datasets was processed twice: first, with the full reference panel of 90 HPRC haplotypes; and then in a leave-one-out (LOO) setting, where the two relevant sample haplotypes were removed from the database beforehand.

To measure genotyping error, we calculated sequence divergence between actual and predicted haplotypes (Figure 2a) and corresponding Phred-like ^24^ quality values (QV), widely used for genome assembly evaluation ^25,26^. Then, we distributed haplotype predictions into four bins based on their QV (<17, 17–23, 23–33 and ≥33), where a haplotype from the last bin (QV ≥ 33) differs from the actual haplotype by no more than 5 bp per 10 kb (see Figure 2b), which is competitive with long read genome assemblies from ONT data ^27^. After genotyping, we marked potentially incorrect genotype predictions based on the number of unexplained reads and the similarities between top genotype predictions; consequently, we will split genotypes into those that *failed* and those that *passed* filtering. Note that the haplotypes were compared across the whole locus, including both coding and non-coding regions, which avoids the need for gene annotations on highly variable haplotypes.

Using the full reference panel, 20,028 Locityper haplotypes (98.4%) passed filtering out of a total of 20,350 fully assembled sample-locus haplotypes across the 256 CMR loci and 40 Illumina WGS samples. Among passed predictions, 94.9% haplotypes had QV ≥ 33, and additional 3.1% haplotypes had QV between 23 and 33 (see Figures 2c and 3). Although some genes remain challenging for accurate genotyping, at 194 (234) loci ≥ 95% haplotypes had QV ≥ 33 (≥ 23).

Even though HPRC assemblies are very accurate, they may include assembly or phasing errors, especially at challenging loci. To remove this factor from the performance analysis, we used ART Illumina ^28^ to simulate 40 short-read datasets and processed them with Locityper. Unsurprisingly, the tool showed even higher accuracy on simulated datasets, producing 99.1% genotypes, which passed filtering; of them 98.6% with QV ≥ 33 (Figure 2c and Supp. Figure 1a).

### Locityper significantly outperforms state-of-the-art short-read variant calling pipelines

Recently, New York Genome Center (NYGC) researchers presented an extensive variant calling pipeline and used to it to call phased single nucleotide variants, indels and structural variants across high coverage Illumina WGS datasets for 3,202 samples from the expanded 1KGP (1000 Genomes Project) cohort ^3^. As the 1KGP call set is phased, we were able to reconstruct local haplotypes at the 256 CMR loci, as well as calculate genotyping accuracy for 39 HPRC samples present in the 1KGP sample set. Even though the NYGC pipeline utilizes state-of-the-art variant callers, 1KGP haplotypes had significant divergence from the actual sample haplotypes, with only 27.4% haplotypes showing QV ≥ 33 and another 22.3% haplotypes with QV < 17 (see Figure 2c). In total, 1KGP haplotypes were overwhelmingly accurate (≥ 95% haplotypes at QV ≥ 33) at only 22 loci (at 83 loci for QV ≥ 23; see Supp. Figure 1b).

**Figure 2.**
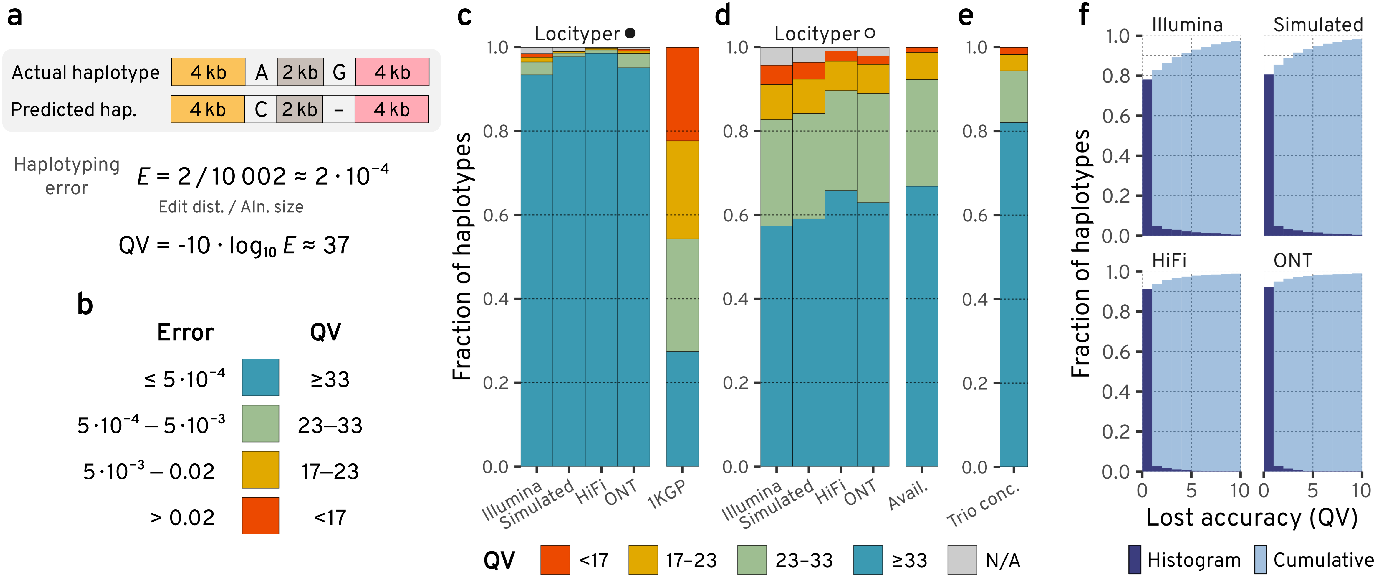
Haplotype accuracy definition and analysis at 256 challenging medically relevant loci. **a**, Haplotyping error is calculated as sequence divergence between actual and predicted haplotypes. Quality value (QV) is a Phred-like transformation of the haplotyping error. **b**, Approximate correspondence between sequence divergence (haplotyping error) and QV bins. **c**, Haplotyping accuracy for full-database Locityper (marked by a filled circle) and the 1KGP call set for up to 40 HPRC samples, respectively. Haplotypes that failed filtering are shown with gray. **d**, Locityper accuracy in the leave-one-out setting (LOO; shown with a white circle) and the corresponding haplotype availability (QV between actual and closest available haplotypes) across up to 40 HPRC samples. **e**, Locityper concordance at 602 Illumina WGS trios from the 1KGP. **f**, Accuracy, lost by Locityper in the LOO setting across HPRC samples—histogram of differences between best available and predicted QVs. Cumulative fraction is shown with light blue.

**Figure 3.**
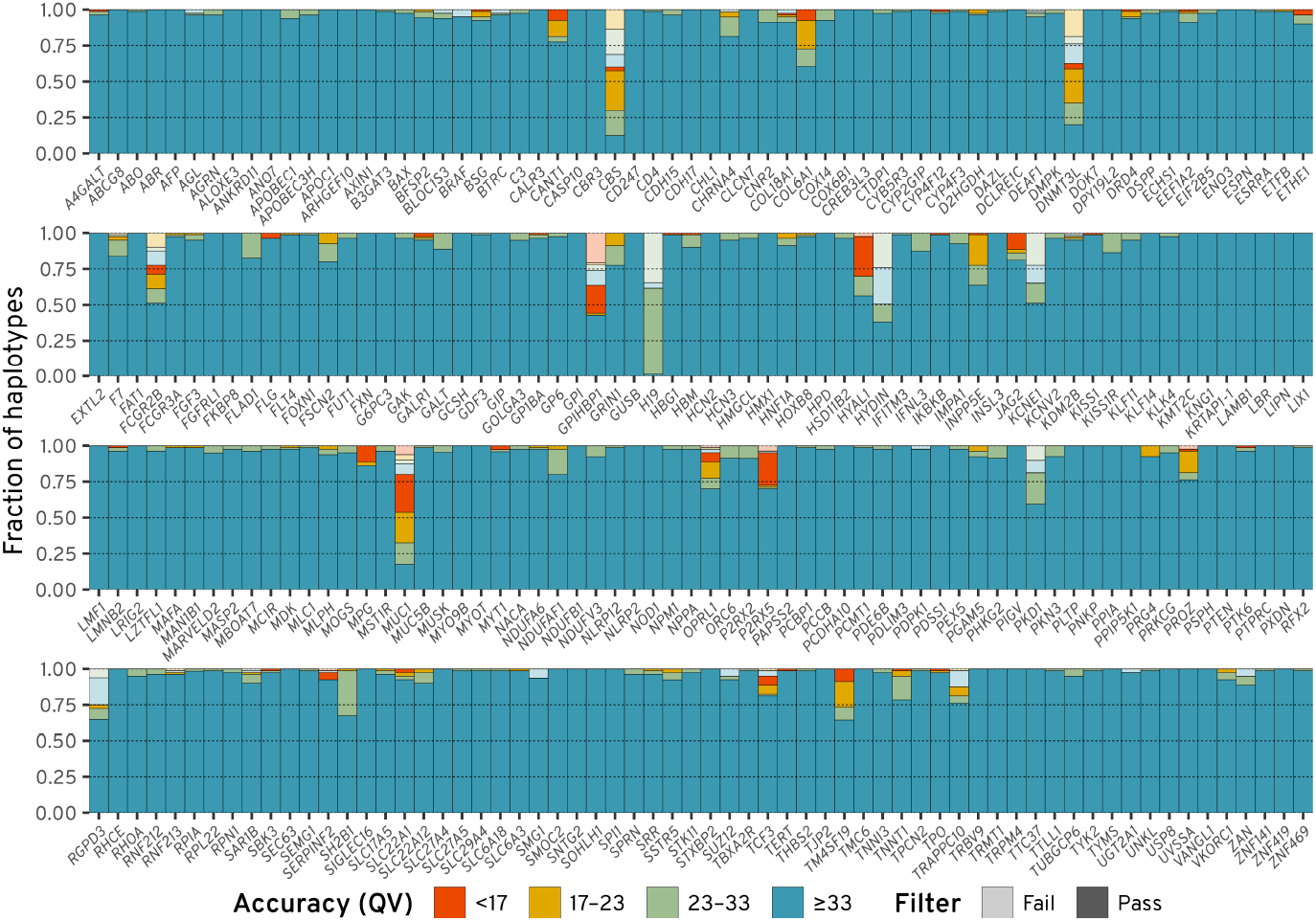
Locityper haplotyping accuracy for 40 Illumina WGS datasets. Predicted haplotypes across 256 challenging medically relevant loci are binned into four groups according to their haplotyping quality value (QV). Semi-transparent colors show predictions that failed post-genotyping filtering.

### Locityper produces high accuracy genotypes based on long reads

Locityper is not limited to short reads: it is able to process various long read WGS datasets, including accurate PacBio HiFi and error-prone Oxford Nanopore (ONT) data (see Figure 2c). For these two technologies, 99.8% and 99.5% Locityper haplotypes passed filtering, respectively; of them, 98.7% and 95.6% were very accurate with QV ≥ 33, while only 0.1% and 0.5% haplotypes had QV < 17, respectively (see Supp. Figure 1c–d).

### Locityper achieves near optimal accuracy in the leave-one-out setting

In the LOO setting, Locityper produced 95.7% Illumina-based haplotypes that passed filtering; among them 86.4% haplotypes had QV ≥ 23, including 59.9% haplotypes with QV over 33 (Figure 2d and Supp. Figure 2a). Even though Locityper achieves smaller accuracy in the LOO setting, it still reconstructs 2.2 times as many highly accurate haplotypes (QV ≥ 33) compared to the 1KGP call set and 4.6 times smaller number of inaccurate haplotypes (QV < 17).

By design, Locityper always associates an input WGS sample with two existing locus haplotypes, and it is not able to predict haplotypes missing from the database. Therefore, Locityper LOO accuracy is limited to haplotype *availability*—similarity between the actual haplotypes and the closest haplotype remaining in the LOO database. 66.8% haplotypes across 40 samples and 256 CMR loci had high availability (closest haplotype with QV ≥ 33); this percentage rises to 92.3% when considering closest haplotypes with QV ≥ 23 (Figure 2d and Supp. Figure 3). In general, Locityper was able to predict haplotypes, close to the best available: *lost accuracy* (difference between best possible and predicted QVs) was under 5 and 10 for 91.4% and 97.3% haplotypes, respectively (see Figure 2f).

At other sequencing datasets, predicted haplotypes were even closer to optimal: lost accuracy was under 5 QV for 93.3%, 97.5% and 97.8% haplotypes based on Simulated, HiFi and ONT reads, respectively; and was under 10 for 98.2%, 99.0% and 99.1% haplotypes (Figure 2f and Supp. Figure 2b– d). In particular, 99.1% HiFi-based LOO haplotypes passed post-genotyping filters and of them 66.4% and 90.5% had QV ≥ 33 and ≥ 23 to the actual haplotypes, respectively.

This analysis shows that Locityper performs extremely well when required haplotypes are present in the reference panel, and achieves near-optimal accuracy with only limited haplotype sets. Growing number of haplotypes in pangenomes ^15^ are likely to increase Locityper accuracy even further.

### Locityper produces concordant trio predictions

In addition to the HPRC dataset, we genotyped the full 1KGP cohort of 3202 Illumina WGS samples, including 602 trios. At each of the 256 CMR loci and for each trio we calculated concordance—similarity between child and parent haplotypes (see Methods). As Figure 2e shows, the vast majority of trio haplotypes were concordant (82.0% and 94.4% with QV ≥ 33 and ≥ 23, respectively). Moreover, average concordance QV surpassed 42 and did not drop below 31 at any locus (see Supp. Figure 4).

### Locityper accurately genotypes HLA and KIR genes

In order to evaluate Locityper ability to genotype hyperpolymorphic genes, we examined genes from two medically relevant genomic regions: Major histocompatibility locus (MHC) covering over 4 Mb and over 200 genes ^29^; and killer-cell immunoglobulin-like receptor (KIR) gene cluster spanning 150 kb and 17 genes ^30^. The two regions contain extremely polymorphic HLA and KIR genes that play an essential role in adaptive and innate immune systems ^31,32^. As Locityper genotypes target loci based solely on the sequences of available haplotypes, it is not limited to gene bodies and can utilize intergenic sequence, gene order and presence/absence of copy-number-variable genes. On the other hand, Locityper may require a large collection of assembled MHC or KIR haplotypes to accurately genotype out-of-sample individuals.

Multiple specialized tools have been developed for genotyping the MHC locus ^19,20,33^; the newest of them being T1K ^21^, a state-of-the-art ^34^ genotyper for HLA and KIR genes that is capable of processing whole genome and whole exome short read sequencing data. To compare T1K and Locityper accuracy, we genotyped 40 Illumina HPRC WGS datasets at 23 genes and 13 pseudo genes from the MHC locus as well as 6 genes and 3 pseudogenes from the KIR locus, all combined into 25 target loci with sum length slightly over 1 Mb. Similarly to CMR loci benchmarking, we run Locityper twice: using the full database and in the LOO configuration.

Across the 40 HPRC samples and 36 genes from the MHC locus, Locityper achieved full match with baseline annotation (correctly predicted all fields in the HLA nomenclature ^35,36^) in 99.5% cases with the full database and 87.9% in the LOO setting, compared to T1K’s 65.0% (Figure 4a,c). When not requiring full matches and evaluating copy-number accurate protein product prediction (second nomenclature field), T1K accuracy rose to 78.3%, while Locityper accuracy grew to 99.5% and 94.0% using full and LOO databases, respectively. Similarly, at the 9 KIR genes, Locityper correctly predicted protein products in 99.9% (full database) and 73.3% (LOO) and achieved full match in 99.7% and 67.9% cases, respectively. Using the same metrics, T1K accuracy reached 68.0% (protein product) and 55.9% (full match) (see Figure 4b–c).

**Figure 4.**
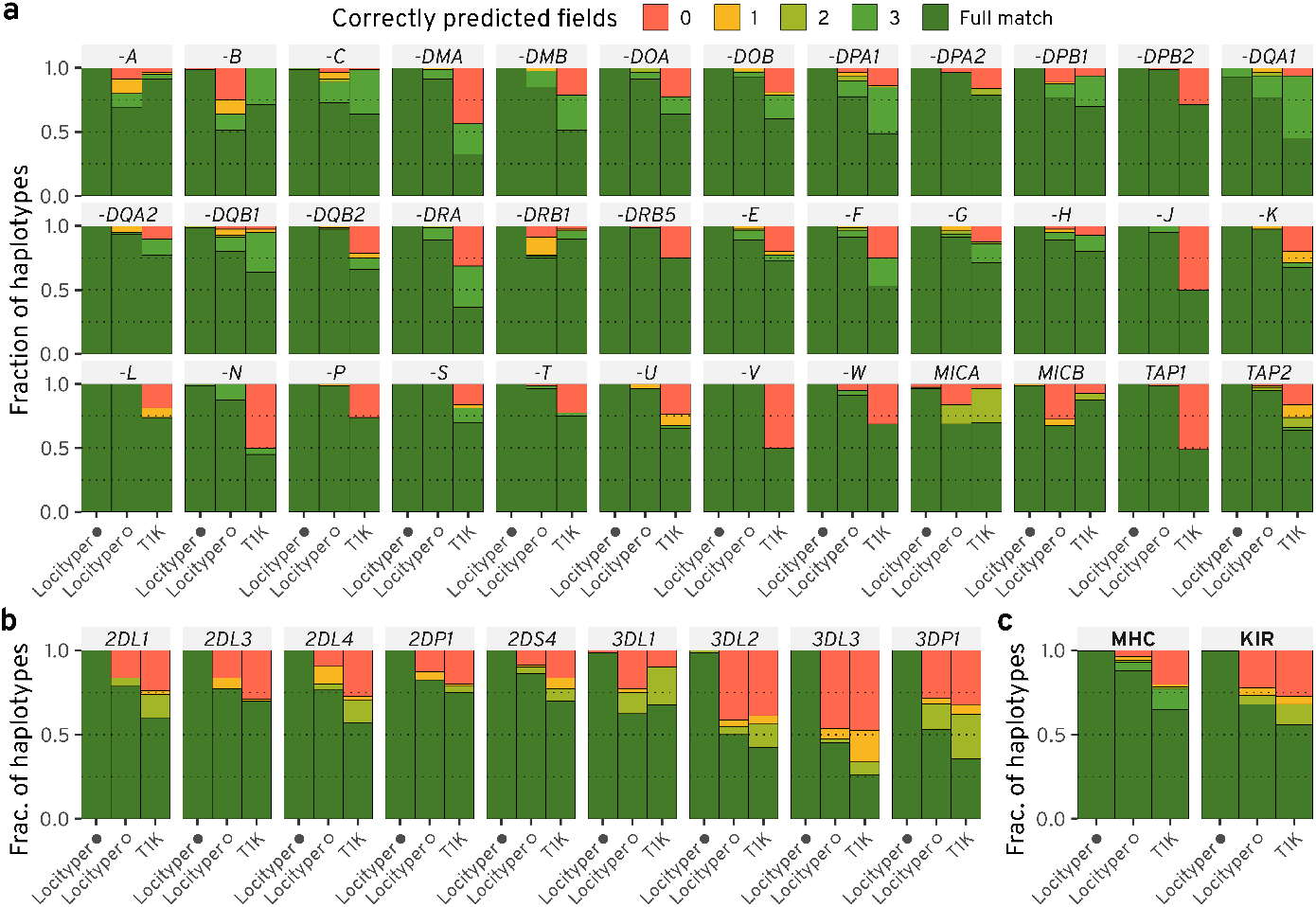
Haplotyping accuracy for 40 HPRC samples at the MHC and KIR loci. Accuracy is shown for Locityper with the full database (denoted by black circle); Locityper in the leave-one-out setting (white circle); and T1K. Fully predicted alleles, as well as correctly identified missing copies, are colored with dark green *(Full match)*, due to the varying number of allele fields in the HLA/KIR gene nomenclature ^35,37^. Otherwise, haplotypes are colored according to the number of correctly predicted fields. **a**–**b**, Haplotyping accuracy at 36 (pseudo)genes from the MHC locus (**a**) and 9 (pseudo)genes from the KIR gene cluster (**b**). **c**, Fraction of haplotypes of various accuracy, aggregated across all MHC and KIR genes/pseudogenes.

Some protein products were present in only one HPRC sample, consequently, such samples cannot be correctly annotated by Locityper in the LOO setting. Cases like that appeared in the most polymorphic HLA and KIR genes and explained 41.3% and 19.0% Locityper LOO errors, respectively (see Supp. Figure 5). At the same time, in many cases T1K predicted smaller copy number than required, explaining 85.7% and 42.5% of all errors at the MHC and KIR loci, respectively. When ignoring these two error types (missing allele predictions and unavailable protein groups), Locityper and T1K achieve similar accuracy when predicting protein products—96.4% and 96.2% at the MHC locus, as well as 77.5% and 78.7% at the KIR genes (Supp. Figure 5). Overall, the general purpose tool Locityper performs in a competitive manner even when compared to T1K, which is specifically designed for HLA/KIR genes.

### Locityper accurately genotypes disease associated gene families

Although the set of CMR genes includes a wide variety of genetically diverse genes, several important polymorphic gene families are underrepresented in it. One such highly heterogeneous gene family is the mucin gene cluster (*MUC1*–*MUC24*) ^40^. Mucin genes encode large glycoproteins that are essential to barrier maintenance and the defense of epithelial tissues. All canonical mucins harbor a large exon that contains Variable Number Tandem Repeats (VNTRs). These VNTR sequences vary per mucin, yet each extensively encode serine and threonine residues for glycosylation ^41^. The gene family can be broken up into two subgroups—that of the cell surface, i.e. tethered mucins and the secreted mucins. In the tethered mucins, single VNTR domains contain variation in total motif copy number and motif usage (Figure 5a); however, the secreted, gel-forming mucins harbor potential variation in VNTR domain copy number, VNTR motif copy number, VNTR motif usage, and cys domain copy number ^42,43^ (Figure 5b).

**Figure 5.**
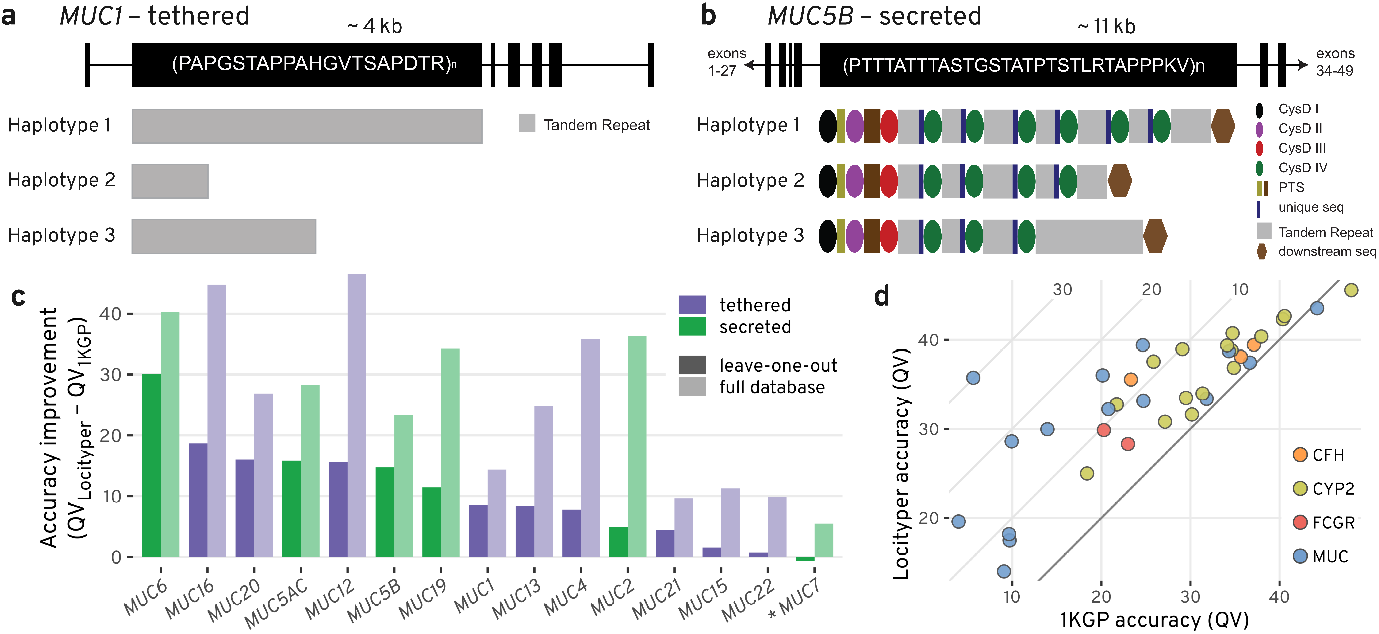
Locityper can accurately genotype mucin and other gene families. **a**, Gene model of *MUC1*, a mucin tethered to the surface of epithelial cells. *MUC1* harbors a 20 amino acid VNTR repeat sequence and is highly polymorphic in VNTR length, as represented by the example haplotypes 1–3^38^. **b**, Gene model of *MUC5B*, a secreted, gel-forming mucin that is important for homeostasis in the lungs. *MUC5B* encodes an irregular 29 amino-acid VNTR motif that is broken up into separate VNTR domains by cys domains. The number of VNTR domains, cys domains, and VNTR motifs could each contribute to polymorphism among haplotypes at this locus ^39^. **c**, Difference in average haplotyping accuracy (QV) between Locityper and 1KGP call set at 15 mucin genes based on 39 Illumina WGS datasets. Improvement for the leave-one-out setting and the full Locityper database are shown with dark and light shades, respectively. Tethered and secreted mucins are shown with purple and green colors; the only non-gel-forming secreted mucin *MUC7* is marked with an asterisk. **d**, Locityper (LOO) and 1KGP call set average genotyping accuracy (QV) across 4 gene families: CFH in orange; CYP2 in light green; FCGR in red; and MUC in blue. Diagonal black line shows zero improvement boundary and diagonal gray lines show 10, 20 and 30 QV improvement.

The presence of these repetitive sequences makes mucins both highly polymorphic and difficult to accurately sequence/genotype in short reads.

Locityper leverages information about both read depth and read alignment for genotyping; therefore, the tool is well suited for the characterization of mucin genetic variation. Based on 39 HPRC Illumina WGS datasets, Locityper (LOO) haplotypes achieved on average 10.5 higher QV compared to the 1KGP call set across 15 examined MUC loci, with the largest improvement observed at *MUC6* and *MUC16* with 30.1 and 18.7 QV, respectively (Figure 5c). The only negative QV difference between Locityper and 1KGP was observed at the non-gel forming *MUC7* gene, where the two haplotype sets showed very high QV values 43.5 and 44.2, respectively.

Further examples of genes that are challenging to address with standard calling techniques are *FCGR2B* and *FCGR3A*, encoding receptors for the Fc region of the immunoglobulin gamma complexes (IgG) ^44,45^. IgG binding to FCGR2B induces immune complexes phagocytosis/endocytosis and therewith establishes the basis of antibody production by B-cells. Several transcript variants encoding different isoforms with differing biological function are present (e.g. isoform IIB2 does not trigger phagocytosis unlike other isoforms ^46^). Genetic variations in this gene have been reported to cause increased susceptibility to systemic lupus erythematosus (SLE) ^47^, as well as to malaria, autoimmune hypersensitivity disease, and immune thrombocytopenic purpura ^48–50^. The second receptor, FCGR3A, is expressed on natural killer cells (NKc) as an integral membrane glycoprotein ^45^. It binds antigen-IgG complexes formed during infection events and thus triggers cytokine production and degranulation by the NK cells. This process is a central effector mechanism limiting viral load and viral propagation in a memory-like manner ^51^. Similar to *FCGR2B*, genetic variants in the *FCGR3A* gene have been associated with SLE ^47^, as well as immunodeficiency, susceptibility to recurrent viral infections, and alloimmune neonatal neutropenia ^52–56^. However, genetic analyses of the FCGR genes using high resolution short reads have been proven notoriously difficult due to a recent gene duplication and diversification processes ^57^. Nevertheless, at the *FCGR2B* and *FCGR3A* receptor genes Locityper (LOO) improves average QV by 5.3 and 9.6 points compared to the 1KGP call set, respectively (23.0 → 28.3 and 20.3 → 29.9) based on the 39 Illumina WGS datasets (Figure 5d). A larger reference panel would likely improve Locityper’s ability to genotype FCGR genes even further, as the tool achieves much higher accuracy (35.9 and 54.0) when using its full database.

Moreover, Locityper (LOO) achieves significant QV improvement (12.2) at the *CFH* gene, associated with age-related vision loss and kidney disorders ^58,59^. Finally, Locityper showed on average 4.5 higher QV across 16 protein coding CYP2 genes that play a major role in drug metabolism ^60,61^. Out of the CYP2 genes, Locityper achieved the highest improvement at *CYP2U1* (9.8), *CYP2A13* (11.0) and *CYP2W1* (11.7) (see Figure 5d).

### Locityper produces more accurate variant calls compared to Pangenie and 1KGP call sets

Pangenie is a pangenome-based short read variant caller, which calls sequence variants and structural variations by counting read *k*-mers along paths in a pangenomic graph ^18^. In order to facilitate comparison to unphased Pangenie call sets, we converted Locityper–predicted locus haplotypes into phased diploid variant calls, and compared Pangenie and Locityper call sets to the ground truth variant calls, extracted from the phased whole genome assemblies for the 40 HPRC samples. Both Locityper and Pangenie were run using their full respective reference panels, which include HPRC samples.

Across the 256 CMR loci, Locityper achieves higher average *F*_1_ score (0.939, median = 0.965) than Pangenie (0.842, median = 0.902), improving both average precision (Pangenie: 0.848, Locityper: 0.945); and recall (Pangenie: 0.843, Locityper: 0.936) by over 9% (see Supp. Figure 6). Although 1KGP call set shows similarly high precision (0.946), it does so at the expense of much lower recall (0.658), achieving a combined *F*_1_ score of 0.746 (median = 0.829). The difference is more pronounced at indels and structural variations, where Locityper, Pangenie and 1KGP call sets attain average *F*_1_ scores of 0.882, 0.728 and 0.550, respectively.

### Runtime and memory usage

Locityper WGS preprocessing (executed once per dataset) took on average 16m (minutes) using 8 threads and consumed 15 Gb of RAM for 30× unmapped Illumina WGS input dataset. If a dataset with a similar library preparation was previously preprocessed, read mapping can be skipped, which speeds up WGS preprocessing to under 3m. Next Locityper step, read recruitment, can simultaneously identify reads for multiple target loci. Due to the fact that reading and decompressing input data was the most time-consuming operation, recruitment speed did not depend on the number of loci (1 to 256 tested). Using 8 threads, read recruitment lasted 14m and had negligible memory footprint (<1 Gb). The final Locityper step, genotyping, consumed <10 Gb of RAM and required approximately 9 seconds per locus, depending the target complexity and size: in total, genotyping 256 CMR loci took 38m30s, while genotyping 25 loci covering MHC and KIR loci took 3m55s. Altogether, Locityper analysis of the HLA and KIR genes, including preprocessing and read recruitment, required under 35 minutes using 8 threads.

At the same time, T1K with 8 threads required on average 2h30m and 48m to process the MHC and KIR loci, respectively, and required 2.5 Gb memory. Pangenie calls variants across the whole genome, and, consequently, had heavier runtime and memory footprint: at 24 threads, its pangenome indexing (executed once) and genotyping steps took 34m and 1h40m, respectively, and consumed 60 and 37 Gb of RAM.

In addition to unmapped data, Locityper and T1K can efficiently utilize mapped reads (in BAM/CRAM formats for Locityper and BAM format for T1K) by only recruiting reads aligned to the regions of interest or to alternative contigs, as well as unmapped reads. Additionally, by examining existing alignments, Locityper is able to preprocess WGS datasets almost immediately. Overall, this decreases T1K runtime to 45m and 23m for HLA and KIR loci, respectively, as well as speeds up the full Locityper pipeline for these genes to under 6m.

## Discussion

In this study, we present Locityper, a targeted method for genotyping complex polymorphic genes using both short and long-read whole genome sequencing. Locityper implements fast read recruitment to a collection of target loci, and utilizes a carefully balanced probabilistic model to calculate genotype likelihoods based on read alignment, insert size and read depth profiles. Locityper employs integer linear programming and stochastic optimization to find the most likely genotype for each target locus. Locityper departs from the prevalent variant-centric approach, which we argue constitutes a particular limitation for highly polymorphic loci. In contrast, our approach leverages collections of known haplotype sequences, which can be extracted from a pangenome reference or directly provided by the user. By examining larger regions around genes of interest, Locityper inherently makes use of any available information, including intergenic sequence, gene order, structural variants, and copy number of short tandem repeats. Locityper is easy to install via docker, singularity or conda, only requires easy-to-obtain input files, has a small memory footprint and significantly shorter runtime than both T1K and Pangenie.

We demonstrated Locityper’s accuracy through excellent agreement to both phased genome assemblies and Mendelian consistency across the 602 family trios included in the 1KGP cohort. When evaluated across a wide range of challenging disease-associated genes, Locityper produces significantly more accurate haplotype predictions compared to a state-of-the art phased variant call set on the 1KGP cohort and outperformed genome-wide pangenome-based genome inference using Pangenie. Locityper’s accuracy stays consistently high across various input sequencing technologies, performing well at Illumina, simulated short reads, PacBio HiFi and Oxford Nanopore Technologies (ONT) datasets.

At present, the size of the available collections of reference haplotypes still poses a limitation. To quantify this effect, we performed leave-one-out evaluations, showing that the best available haplotype does not reach QV 33 in more than 30% of cases (Fig. 2d). Therefore, despite Locityper’s ability to predict haplotypes close to the best available, the resulting accuracy is not yet ideal for all genes of interest. Significantly larger pangenomes are presently being constructed by the HPRC ^15^ and we are confident that these future pangenomes will lead to a significant increase in performance on out-of-sample individuals for more complex polymorphic genes. Already now, Locityper outperforms the specialized genotyper T1K across HLA and KIR genes in a LOO setting and shows improved ability to genotype other medically relevant gene families (e.g. MUC and FCGR) using short read WGS.

As part of this study, we used Locityper to process 3,202 Illumina WGS datasets from the 1KGP and make the obtained genotypes available, which provides a resource for deeper analyses of >300 challenging target loci. Additionally, publicly available Locityper-preprocessed WGS summaries will allow for a faster genotyping of genes that were not a focus of this study across the 1KGP cohort. We envision that Locityper will enable the inclusion of complex loci in GWAS ^62^ and PheWAS ^63^ analyses, especially in a larger cohort, such as the All-of-Us program ^22^ and the UK Biobank ^23^, which promises to discover many new associations and explain missing heritability. Of note, Locityper’s ability to process both short and long reads might prove especially useful for the increasing production of long reads in the context of biobank-scale sequencing efforts.

For a given locus, Locityper aims to find two existing haplotypes that would explain an input WGS dataset in the best way. Consequently, it is not designed to reconstruct a novel haplotype, even if it constitutes a mixture of already known haplotypes. To address this, Locityper outputs read alignments to the top predicted genotypes, which can be used later for visual analysis or variant calling. Combined with assembly polishing ^64,65^, this could improve genotyping accuracy and allow for reconstruction of previously unobserved alleles—a strategy that we plan to explore in future research.

Different parts of a gene of interest, such as exons, introns, and tandem repeats, might have different relative impact on its biological function, as well as on the allele/protein groups in the corresponding allele classification. Currently, Locityper weights all sequence windows according to their *k*-mer content and sequence complexity. However, a modification can be made that would either automatically, or with user input, upweight/downweight various parts of the haplotype according to their phenotypic importance. Moreover, two loci with significant homology, e.g. part of a non-tandem segmental duplication, are processed independently, with potentially overlapping sets of recruited reads. Locityper mitigates this problem by tracking the number of off-target *k*-mers per read/haplotype window. Nevertheless, further method improvements are conceivable, such as using a shared pool of reads for the related loci, similar to the strategy implemented by T1K ^21^.

In conclusion, Locityper allows for fast and accurate targeted genotyping of challenging polymorphic loci using various sequencing technologies. With the current draft pangenome containing highly accurate phased-genome assemblies, Locityper routinely achieves sequence accuracies above QV 33, which is comparable to genome assemblies from Oxford Nanopore data ^27^. As more human haplotypes are represented in pangenomes, we expect the accuracy to improve further, which will facilitate detailed analysis of previously intractable genes, leading to improved diagnostic power and novel disease associations.

## Data availability

Locityper-predicted genotypes for 3202 Illumina 1KGP samples, corresponding preprocessed WGS parameters, target loci database and benchmarking results can be found on Zenodo ^66^ (https://doi.org/10.5281/zenodo.10977559).

## Code availability

Locityper is implemented in the Rust programming language, and can be installed via conda, singularity and docker. Source code is freely available under the terms of the MIT license at github.com/tprodanov/locityper along with installation and usage instructions. Fixed source code version as well as additional benchmarking scripts can be downloaded from Zenodo ^67^ (doi.org/10.5281/zenodo.10979046).

## Methods

In this article, we present a targeted tool *Locityper*, designed for genotyping complex multi-allelic loci. Locityper processes whole genome sequencing (WGS) data produced by various sequencing technologies, including highly accurate short and long reads (such as Illumina and PacBio HiFi data, respectively), as well as error-prone long reads (such as PacBio CLR and Oxford Nanopore data). Locityper can efficiently analyze unmapped reads stored in various formats, as well as mapped reads from sorted and indexed BAM/CRAM files.

Broadly, the method can be split into several steps:

1. Preprocessing target loci,
2. Sample preprocessing (performed once for each WGS dataset),
3. Read recruitment (carried out simultaneously for multiple loci),
4. Locus genotyping and generating BAM files with alignments to the best genotypes. These steps are described in more detail in the following sections.

### Preprocessing target loci

Locityper utilizes solely locus haplotype sequences, and does not require any kind of graph structure on top of them. Locus haplotypes can be provided directly in a FASTA file; alternatively, Locityper can automatically extract locus haplotypes from a pangenome, provided in a VCF format (constructed, for example, by Minigraph-Cactus ^68^).

When locus haplotypes are extracted from a VCF file, Locityper tries to extend the locus in such a way that both locus ends do not overlap any pangenomic variation. Additionally, the tool tries to select a position that would produce the largest number of unique canonical *k*-mers at the edges of the locus (default edge size = 500 bp). In the default configuration, locus extension is limited by 50 kb at each side, but can fail if there is a longer structural variant at the locus boundary. In such cases, the user can either increase the allowed extension size, or set the boundaries manually.

Next, Locityper removes all identical haplotypes and calculates Jaccard distance ^69^ between multisets of minimizers ^70^ ((15, 15) by default) for all pairs of haplotypes. These pairwise distances are later used to flag potentially incorrect predicted genotypes. Finally, Locityper finds off-target *k*-mer multiplicities, calculated as the difference between canonical *k*-mer counts across the full reference genome (calculated using Jellyfish ^71^; recommended *k* = 25) and the corresponding *k*-mer counts at the reference locus sequence.

### WGS dataset preprocessing

Locityper aims to probabilistically describe three features of a given WGS dataset— insert size, error profile and read depth— by examining read alignments to a predefined background region. For human WGS data, we use a 4.5 Mb interval on the chr17q25.1 as the default background region as it contains almost no segmental duplications or other types of structural variations. If input reads are unmapped, Locityper subsamples input reads by a factor of *s* (^1^/_10_ by default) and maps them to the reference genome using Strobealign ^72^ (short reads) or Minimap2^73^ (long reads).

#### Insert size

Manual examination of several paired-end WGS datasets from the HPRC project ^15^ indicated that the Negative Binomial (NB) distribution fits insert size distribution the best (see Supp. Figure 7). For a given WGS dataset, we use all fully mapped read pairs (clipping < 2% of the read length, by default) with high mapping quality (≥ 20). We remove outliers by defining maximum allowed insert size as three times the 99^th^ percentile of the observed insert sizes, and discard violating read pairs. Finally, we obtain the NB distribution parameters using the method of moments. During the next two preprocessing steps we will only use read pairs with insert sizes within the 99.9% confidence interval of the corresponding NB distribution.

#### Error profile

We use two distributions to describe WGS error profiles. First, we use Beta-Binomial (BB) distribution to evaluate edit distance based on the read length. The distribution is fitted using the maximum likelihood estimation (MLE) based on the remaining read pairs. Obtained BB distribution will be used to distinguish between true and off-target alignments at the genotyping stage.

Second, we calculate match, mismatch, insertion and deletion rates (*p*_*M*_ , *p*_*X*_ , *p*_*I*_ , *p*_*D*_, respectively), and define alignment likelihood as the product of the corresponding rates to the power of the number of operations. For example, alignment with 100 matches, 1 mismatch and 2 insertions would receive likelihood 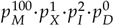. Note that the probabilities do not sum up to one and are incomparable between reads of different lengths. Nevertheless, this formulation produces fast to calculate probabilities, and provides a way to numerically compare different alignments of the same read.

#### Read depth

We split the background region into windows of fixed size based on the mean read length, and assign reads to windows based on the middle of the corresponding read alignments. Next, we count the number of primary read alignments assigned to each window (only first mates are counted to preserve window independence) ^74^.

For each window we calculate GC-content and the fraction of unique *k*-mers in an area centered around the window. Next, we select windows with many unique *k*-mers (≥ 90%) and estimate read depth mean and variance across various GC-content values using local polynomial regression ^75^. NB parameters are then estimated separately for each GC-content based on the smoothed mean, variance, and subsampling rate (see Supplementary Methods 3.1).

### Read recruitment

Following dataset preprocessing, Locityper recruits reads to all target loci. For that, we collect minimizers ^70^ from each locus and each haplotype (default: (5,15) and (10,15)-minimizers for short and long reads, respectively). Uninformative minimizers, which appear ≥ 5 times off target, are ignored. Locityper compares read and target minimizers in parallel and recruits reads to one or several loci according to one of the following rules: short reads are recruited if a sufficient fraction of minimizers matches the target for all read ends (default: 0.7 and 0.6 for single- and paired-reads).

Only a small part of a long read may overlap a given target locus. Consequently, we recruit a long read if it contains a subregion with sufficiently many minimizer matches. For that, we employ the following heuristic: matching/mismatching informative minimizers are assigned *s*_+_/*s*_−_ scores (default: +3/-1), and a read is recruited if it has a continuous subsequence with sum score greater or equal to

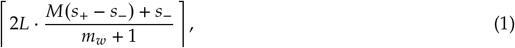

where *L* is the subregion length (default: 2000 bp), *M* is the match fraction (default: 0.5) and 2*L*/(*m*_*w*_ + 1) is the expected number of (*m*_*w*_ , *m*_*k*_ )-minimizers per *L* bp sequence ^76^. This heuristic is useful as it can be quickly evaluated using Kadane’s algorithm ^77^, and as it is not too restrictive: shorter read subregions with a higher match rate may produce a hit, and vice versa.

### Genotype likelihood

#### Read location probabilities

Following the read recruitment, every target locus is genotyped independently from other loci. Reads, recruited to the locus, are aligned to all haplotypes *H* using either Strobealign ^72^ or Minimap2^73^, depending on the read type. Obtained read alignments are assigned BB *p*-values according to their edit distances and read lengths. A read pair is retained if both read ends have at least one good alignment (*p* ≥ 0.01) to at least one of the haplotypes. All alignments with BB *p* < 0.001 are discarded.

Without loss of generality, we will describe the following steps for paired-end reads and use notation **r** = (*r*_1_, *r*_2_) to describe a read pair. Each locus haplotype *h* ∈ *H* is split into non-overlapping windows *W*^(*h*)^ of fixed size (same size as in read depth preprocessing); furthermore, we expand *W*^(*h*)^ by adding a null window *w*_○_. Each alignment is connected to a single window *w* based on the middle point of the alignment, with alignment probability 𝔭(*r*_*j*_ , *w*) calculated according to the precomputed error profile. Reads without proper alignment to *h* are connected to the null window *w*_○_; we will define 𝔭(*r*_*j*_ , *w*_○_) as Λ · max_*h*_ 𝔭(*r*_*j*_ , *h*) — the probability of the best *r*_*j*_ alignment to any haplotype, multiplied by a penalty Λ (10^−5^ by default).

Paired end alignment probability of the read pair **r** = (*r*_1_, *r*_2_) to windows **w** = (*w*_1_, *w*_2_) can be written as 𝔭(**r, w**) = 𝔭(*r*_1_, *w*_1_) · 𝔭(*r*_2_, *w*_2_) · *P*_insert_(**r, w**), where the last term is calculated according to the precomputed insert size distribution. For null windows, we define insert size probability as the highest probability achievable under the precomputed insert size distribution. Finally, we will denote the full set of possible read pair locations on haplotype *h* as 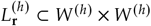 and will define the probability of the read pair **r** location to be **w** as normalized alignment probability:

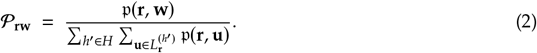

Some parts of the target loci can have high homology to other genomic regions. Consequently, we downgrade the effect of potentially misrecruited reads by setting equal probabilities to all locations for read pairs with less than 5 target-specific *k*-mers.

#### Read assignment

Without loss of generality, let us consider a diploid genotype **g** = (*h*_1_, *h*_2_). We combine windows across the two haplotypes 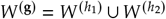. If *h*_1_ = *h*_2_, we use two copies of each window, such that |*W*^(**g**)^| is always 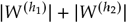. Next, for each read pair **r** we concatenate possible locations 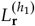 and 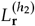 in a similar way to achieve a combined list of locations 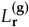.

We describe read assignment to one of the locations using a boolean vector *T*_**r**_ with exactly one true element (Σ _**w**_ *T*_**rw**_ = 1 ), where *T*_**rw**_ = 1 encodes the statement “true location of the read pair **r** is **w**”. Probability of the read assignment *T* for all read pairs *R* is a product of all selected location probabilities:

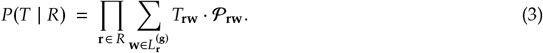

In order to evaluate genotype concordance with WGS data, we search for a read assignment *T* that produces maximum joint probability *P*(**g**, *T* | *R*) = *P*(**g** | *T* , *R*) · *P*(*T* | *R*) = *P*(**g** | *T*) · *P*(*T* | *R*).

#### Read depth likelihood

We define genotype likelihood conditional on the read assignment as the probability for all genotype windows to have copy number (CN) equal to one:

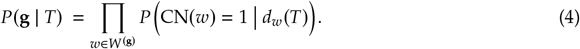

*d*_*w*_(*T*) denotes window *w* depth according to the read assignment *T*, which can be written as 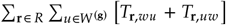. At CN = 1, read depth follows the NB distribution with precomputed parameters *n* and *ψ*. Bayes’ theorem with equal priors produces the following result:

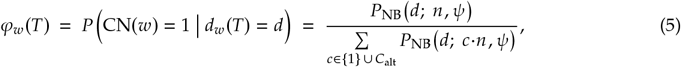

where alternative hypotheses are represented by a set *C*_alt_. We found it beneficial to use *C*_alt_ = {0.5, 1.5}, in other words, a half divergence from the expected read depth is considered significant. As unmapped reads are already penalized by low alignment probabilities 𝔭(*r* , *w*_○_), we define *P* CN(*w*_○_) = 1 *d* = 1 for any read depth *d*.

#### Window and read weights

Low-complexity regions, as well as short and long repeats evoke various difficulties in read sequencing, recruitment and alignment. In order to assign window weights in a continuous fashion, we define the following two-parametric function *ϑ* : [0, 1] ↦→ [0, 1]:

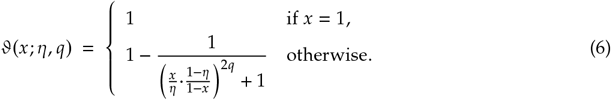

*ϑ* exhibits several useful properties: it is a strictly increasing smooth function such that *ϑ*(0) = 0 and *ϑ*(1) = 1. Location parameter *η* ∈ (0, 1) defines the break point *ϑ*(*η*; *η*, ·) = 1/2, while the power parameter *q* > 0 controls the “slope” of the function, with larger *q* producing larger derivative *ϑ*^′^(*η*; *η, q*) (see Supp. Figure 8). Finally, we define window *w* weight *ζ*_*w*_ = *ϑ*(*x*_1_; *η*_1_, *q*_1_) · *ϑ*(*x*_2_; *η*_2_, *q*_2_) based on the fraction of locus-specific *k*-mers *x*_1_ and linguistic sequence complexity *x*_2_ = *U*_1_*U*_2_*U*_3_, where *U*_*i*_ is the fraction of unique *i*-mers in the window *w* out of the maximal possible number of distinct *i*-mers ^78^, with default parameters *η*_1_ = 0.2, *η*_2_ = 0.5 and *q*_1_ = *q*_2_ = 2.

#### Combined likelihood and likelihood update

Not accounting for window weights, combined likelihood for a genotype **g** and read assignment *T* can be calculated as:

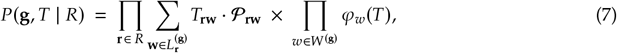

Next, we move the calculations to log-space, add window weights *ζ*_*w*_, and introduce contribution factors Ω_*R*_, Ω_*D*_ ≥ 0, which represent the relative importance of read alignment and read depth likelihoods, respectively. Then, log-likelihood ℒ can be written in the following way:

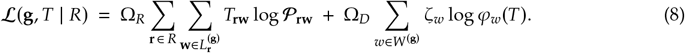

Contribution factors Ω_*R*_ and Ω_*D*_ are necessary as the read alignments can overshadow read depth due to the large number of read pairs and large differences between various read alignments. The two values need to be defined in advance and should sum up to 2: we recommend default values Ω_*R*_ = 0.15, Ω_*D*_ = 1.85, as they produced good results across a selection of target loci and sequencing datasets.

#### Likelihood update

Given log-likelihood ℒ(**g**, *T* | *R*) for genotype **g** and some read assignment *T*, we can efficiently calculate log-likelihood ℒ(**g**, *T*^′^ | *R*) for a new read assignment *T*^′^ if the read assignment has changed for only one read pair; in other words, when 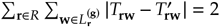. Suppose that the read assignment changed for read pair **r** from location *u, v* (in *T*) to *u*^′^, *v*^′^ (in *T*^′^). Then, read depth likelihood values *φ*_*w*_(*T*^′^) will be identical to *φ*_*w*_(*T*) for all windows except for *u, v, u*^′^, *v*^′^, where read depth can quickly be recomputed. This way, log-likelihood can be recalculated in constant time:

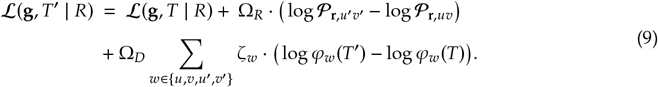

### Finding best read assignment

For each genotype **g** we aim to find such read assignment *T* that would maximize joint log-likelihood ℒ(**g**, *T* | *R*). Locityper implements three approaches for finding such read assignment: Stochastic Greedy approach ^79^, Simulated Annealing ^80^ and Integer Linear Programming (ILP) ^81^. The first two algorithms start from an arbitrarily generated read assignment *T*, then iteratively select a random read pair **r** and switch its location if it increases the genotype likelihood. In addition to “good” location switches, Simulated Annealing permits “bad” switches (decreasing overall likelihood), gradually restricting frequency of such events.

In an ILP formulation we introduce two sets of unknowns: *x*_**rw**_ ∈ {0, 1} for each read pair **r** and each location 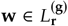; and *y*_*wd*_ ∈ {0, 1} for each window *w* ∈ *W*^(**g**)^ and each possible window depth *d* between zero and maximal possible read depth *D*_max_. The problem can be written as follows:

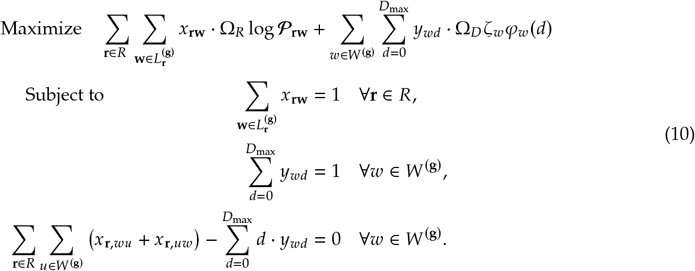

Note, that we can remove variables *x*_**r**_ for trivial read pairs, which map to only one possible location; at the same time, the number of possible read depth variables *y*_*w*_ is exactly one more than the number of non-trivial read pairs mapping to *w*. Finally, the sum 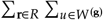 in the third constraint can be limited to windows and read pairs, relevant to the window *w*. Locityper utilizes two commercial ILP solvers, available under academic licenses: HiGHS ^82^ and Gurobi ^83^. Note that it is possible to state a bigger

ILP problem; solution to such formulation would immediately produce the best locus genotype (see Supplementary Methods 3.2). However, we observed that the available ILP solvers are unavailable to quickly and accurately find such solution.

### Locus genotyping

In order to find the best locus genotype for the input WGS data, Locityper finds the best read assignment and the corresponding genotype likelihood for each possible locus genotype (Figure 1). To speed up the process, we start by calculating log-likelihood in the absence of read depth (Ω_*D*_ = 0), which can be efficiently computed by assigning every read to its most probable location. Then, we employ a heuristic filtering by removing all genotypes whose likelihood is 10^100^ smaller than the best likelihood (first 500 genotypes are kept regardless of likelihood). For all remaining genotypes, the best read assignment is found using one of the three approaches, described above. Even though the ILP solvers typically find better read assignments, we use Simulated Annealing as the default solver as it produces decent read assignments in a fraction of ILP solving time.

Splitting locus haplotypes into non-overlapping windows is an intrinsically discrete process. Furthermore, windows can be shifted across different haplotypes due to the presence of indels. Consequently, identical read depth profiles may produce varying read depth likelihoods depending on the window boundaries. To reduce this effect, we perform a procedure, similar to Noise Injection regularization ^84^, where we randomly move read alignment centers to either direction and reassign reads to windows. In addition, we redefine window GC-content values and weights *ζ*_*w*_ as if the window was randomly moved (actual window boundaries stay fixed). In a default configuration, read and window movement is limited to half window size or 200 bp, whichever is smaller. Repeating noise injection several times (20 by default), together with the stochastic nature of likelihood maximization produces a distribution of log-likelihoods for each genotype.

Finally, Locityper selects a *primary* genotype with the highest average log-likelihood and calculates its Phred quality ^24^ based on the probability of error: probability that true log-likelihood of any other genotype is higher than true log-likelihood of the primary genotypes, calculated using one-sided Welch’s *t*-test ^85^.

In addition to quality values, Locityper outputs the number of unexplained reads—reads that map to some, but not to the two predicted haplotypes. Additionally, Locityper iterates over all genotypes and evaluates average Jaccard distance to the primary genotype (see locus preprocessing) weighted by the corresponding genotype probabilities. Such a measure is motivated by the fact that all probable genotypes should be similar to one another. For this study, we marked genotypes as potentially incorrect if weighted distance is over 30 or if there are more than 1000 unexplained reads, which in turn constitute over 20% of all reads for the locus.

### Locus selection

Original set of challenging medically relevant (CMR) genes contains 273 protein coding genes ^7^. We expanded gene coordinates to a minimum of 10 kb, when needed, and supplied positions as input to Locityper locus preprocessing, allowing an additional coordinate expansion by at most 300 kb to each of the sides (add -e 300k). At this stage, eight genes (*ATPAF2, CLIP2, GTF2I, GTF2IRD2, IGHV3-21, MRC1, NCF1* and *SMN1*) were discarded as at least one the gene ends was contained in a 300 kb–long pangenomic bubble. Afterwards, we removed redundant loci (completely contained in another locus), which produced a final set of 256 loci, containing 265 CMR genes.

In a similar fashion, we added 26 loci covering genes from the MHC and KIR gene clusters, as well as 30 loci covering MUC, CFH and CYP2 genes. Full information about the loci can be found in the Supp. Table 1.

### Utilized data

Pangenome reference in a variant calling format (VCF) was downloaded from https://s3-us-west-2.amazonaws.com/human-pangenomics/pangenomes/freeze/freeze1/minigraph-cactus/hprc-v1.1-mc-grch38/hprc-v1.1-mc-grch38.raw.vcf.gz. Illumina, PacBio HiFi and Oxford Nanopore data for the HPRC samples can be found at https://s3-us-west-2.amazonaws.com/human-pangenomics/index.html?prefix=working. NYGC variant calls for the 1KGP samples were downloaded from http://ftp.1000genomes.ebi.ac.uk/vol1/ftp/data_collections/1000G_2504_high_coverage/working/20220422_3202_phased_SNV_INDEL_SV. 3202 1KGP Illumina datasets are available on the European Nucleotide Archive under accession codes PRJEB31736 and PRJEB36890.

Simulated Illumina data was constructed using ART Illumina ^28^ v2.5.8 with parameters -ss HS25 -m 500 -s 20 -l 150 -f 15 for all phased haplotype assemblies from the HPRC project, which can be found at doi.org/10.5281/zenodo.5826274.

### Benchmarking Locityper

In order to evaluate haplotyping accuracy, we computed full-length alignments between actual and predicted haplotypes using Locityper align module. Internally, it finds the longest common subsequence of *k*-mers using LCSk++ ^86^ and completes the alignment between *k*-mer matches using Wavefront alignment algorithm ^87,88^. Three *k*-mer sizes are tried (25, 51 and 101), and an alignment with the highest alignment score is returned.

Afterwards, we calculate haplotyping error—sequence divergence between two haplotypes, calculated as the ratio between edit distance Δ and alignment size *S* (edit distance plus the number of matches). As actual and predicted genotypes consist of two haplotypes, there are two possible actual–predicted haplotype pairings. Out of the two options we select such pairing that produces a smaller ratio between sum edit distance and sum alignment size.

Then, we use Phred-like transformation of haplotyping error QV = −10 · log_10_(Δ/*S*) to obtain haplotyping quality values (QV) ^25,26^. However, when two haplotypes are completely identical (Δ = 0), QV becomes infinite, which poses problems for average QV calculation. For that reason, we corrected QV definition:

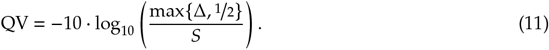

This way, QV difference between edit distances 0 and 1 is the same as between 1 and 2, and equals to 10 · log_10_ 2 ≈ 3. Constants smaller than ^1^/_2_ were generally even more beneficial for Locityper benchmarking. In the leave-one-out setting, we calculate lost accuracy as the difference between best possible QV_avail_ (QV of the closest remaining haplotype) and QV_pred_ of the predicted haplotype. However, as to not penalize well-predicted haplotypes relative to very good possible haplotypes, we modified lost accuracy to be min{QV_avail_, 33} − min{QV_pred_, 33}. This way, QV_pred_ = 30 with QV_avail_ = 50 will produce lost accuracy = 3 instead of 20.

We considered a trio of locus genotypes concordant if one of the child haplotypes matches well one of the maternal haplotypes, while another child haplotype matches one of the paternal haplotypes. Similarly to haplotyping error calculation, we iterated over 8 possible pairings and selected one with the smallest sum edit distance divided by sum alignment size, as well as calculated a QV score for each of the child haplotypes.

We used Bcftools ^89^ (v1.18) consensus command to reconstruct haplotypes from the 1KGP phased variant call set ^3^. In the process, we removed contradicting overlapping variant calls, and variants with symbolic alternative alleles (with exception of <DEL>), as they cannot be used for haplotype reconstruction.

In order to compare Locityper, 1KGP and Pangenie ^18^ v3.02 call sets, we decomposed and normalized variant calls using Vt ^90^ v0.57721 commands decompose_blocksub and normalize, respectively. Before decomposition we removed low quality variants (Locityper genotypes, which failed filtering, and Pangenie variants with quality <10). Then, we used RTG tools ^91^ v3.12.1 vcfeval module to calculate variant calling precision and recall using existing HPRC Minigraph-cactus ^68^ representation as a baseline call set.

Finally, we used T1K ^21^ v1.0.5 with presets hla-wgs --alleleDigitUnits 15 --alleleDelimiter : and kir-wgs with all other parameters set to default. Ground-truth HLA and KIR annotation for HPRC assemblies were obtained using Immuannot ^92^ using allele databases ^31,93^ IPD-IMGT/HLA v3.52 and IPD-KIR v2.12. If a haplotype contains a novel gene allele, Immuannot may associate it with several existing alleles. In such cases, we evaluated predicted allele by the best-matching existing allele.

In all evaluations, we utilized Locityper v0.15.1 along with its dependencies Samtools ^89^ v1.18, Jellyfish ^71^ v2.2.10, Minimap2^73^ v2.26-r1175 and Strobealign ^72^ v0.13.0.

## Acknowledgements

We thank the MHC working group of the Human Genome Structural Variation Consortium (HGSVC) for valuable feedback on an earlier version of the evaluation. This research was supported in part by funding from the National Institutes of Health (NIH) National Human Genome Research Institute (NHGRI) R01 HG002385 (to E.E.E. and T.M.). E.E.E. is an investigator of the Howard Hughes Medical Institute.

## Author contributions

T.P. and T.M. conceived the project and designed the algorithm. T.P. developed the software and performed the analyses. T.P. and E.G.P. prepared the figures. T.P., E.G.P., G.S., S.G.M., E.E.E. and T.M. wrote the manuscript.

## Competing interests

E.E.E. is a scientific advisory board (SAB) member of Variant Bio, Inc.

## Notes

https://github.com/tprodanov/locityper

### Summary of Updates

Reverting to the original manuscript version.

