## Supplementary Information for "Locityper: targeted genotyping of complex polymorphic genes"

### Supplementary Information for
### Locityper: targeted genotyping of complex polymorphic genes

Timofey Prodanov [1,2,✉], Elizabeth G. Plender [3,4], Guiscard Seebohm [5], Sven G. Meuth [6], Evan E. Eichler [3,7] & Tobias Marschall [1,2,✉]

[1] Institute for Medical Biometry and Bioinformatics, Medical Faculty, Heinrich Heine University, 40225 Düsseldorf, Germany. [2] Center for Digital Medicine, Heinrich Heine University, 40225 Düsseldorf, Germany. [3] Department of Genome Sciences, University of Washington School of Medicine, Seattle, WA 98195, USA. [4] Basic Sciences Division and Computational Biology Program, Fred Hutchinson Cancer Center, Seattle, WA 98109, USA. [5] Institute for Genetics of Heart Diseases, Department of Cardiovascular Medicine, University Hospital Münster, 48149 Münster, Germany. [6] Department of Neurology, Medical Faculty, Heinrich Heine University, 40225 Düsseldorf, Germany. [7] Howard Hughes Medical Institute, University of Washington, Seattle, WA 98195, USA.

**1 Supplementary Figures** 1

### 1 Supplementary Figures

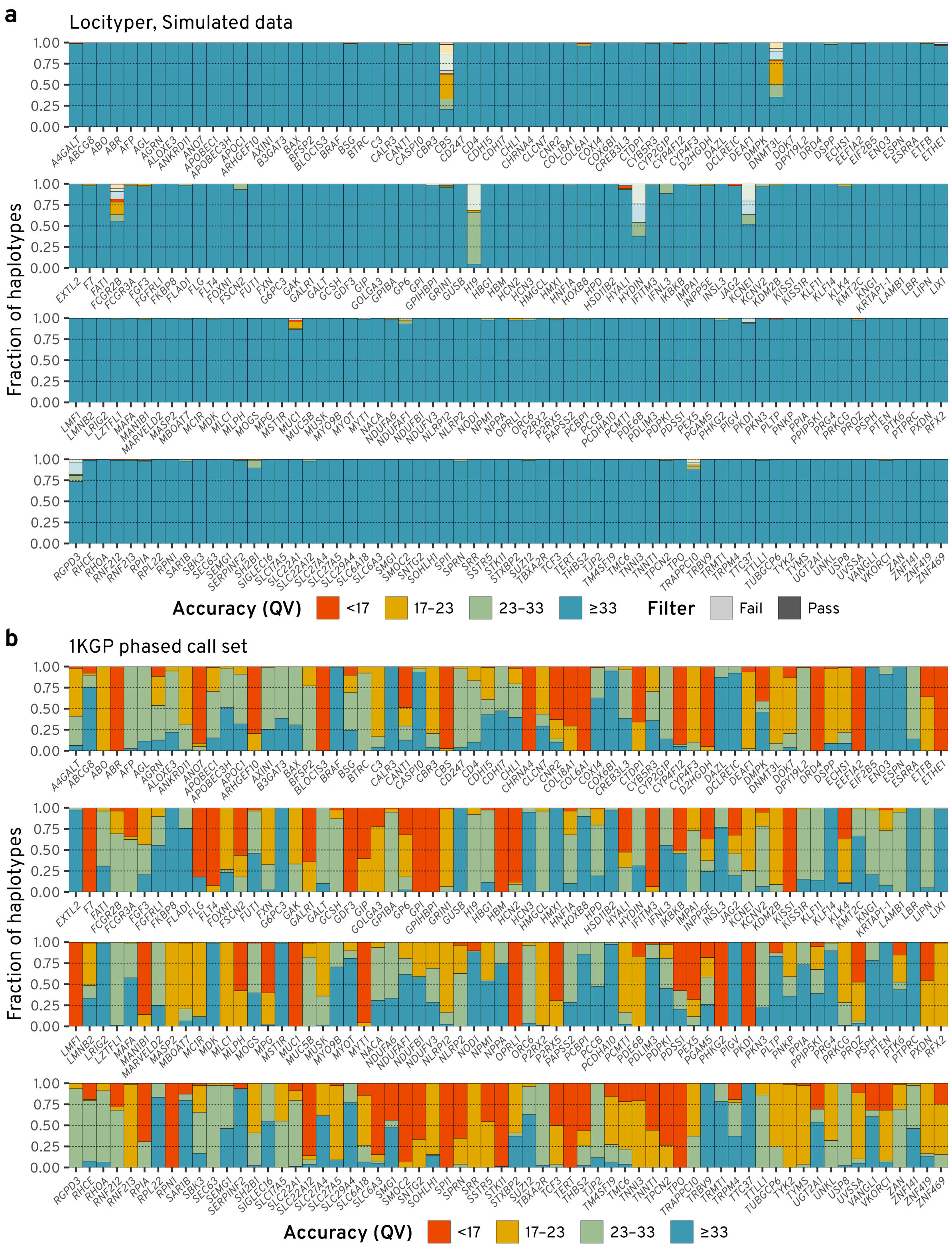

**Supplementary Figure 1. Haplotyping accuracy of Locityper and the 1KGP call set across 256 challenging medically relevant loci.** Haplotypes are stratified into four bins based on their quality values (QV; shown with different colors). Predictions that were discarded during post-genotyping filtering are shown with semi-transparent colors. **a**, Locityper accuracy at 40 simulated short-read datasets. **b**, Accuracy of haplotypes, reconstructed from the phased 1KGP call set for 39 HPRC samples. *[Continued on the next page]*

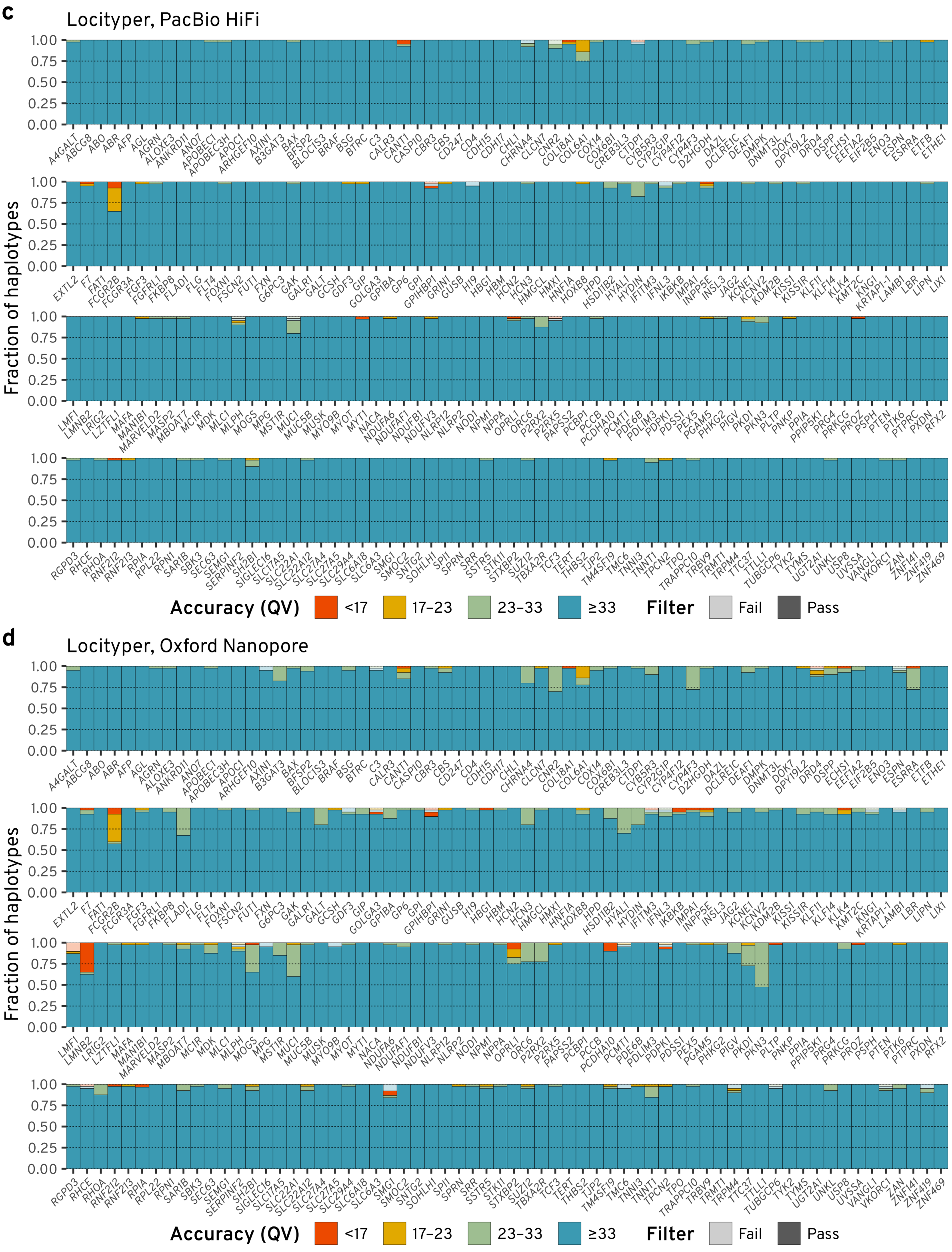

**Supplementary Figure 1.** *(continued)* **c**, Locityper accuracy at 20 PacBio HiFi datasets. **d**, Locityper accuracy at 20 Oxford Nanopore datasets.

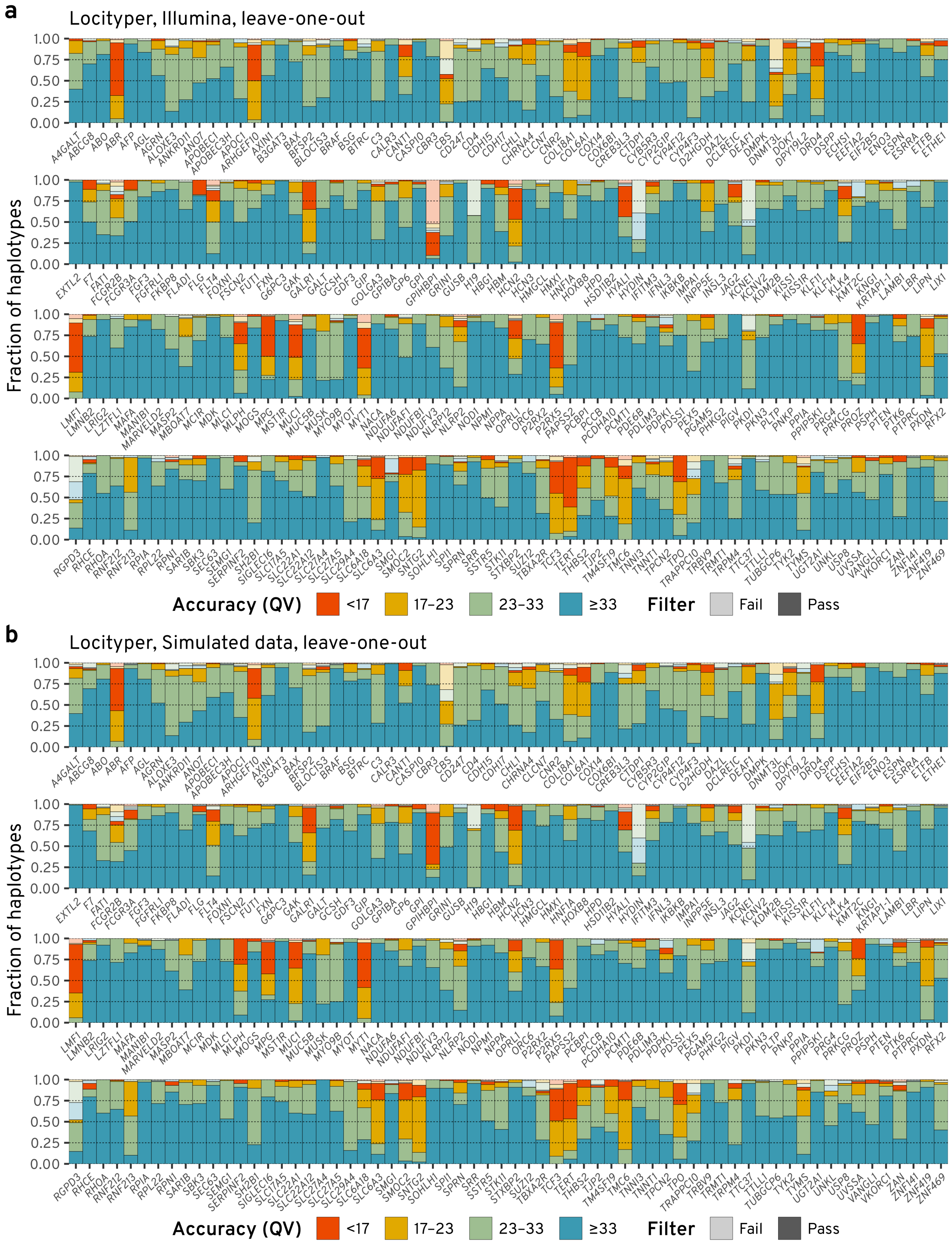

**Supplementary Figure 2. Haplotyping accuracy of Locityper in the leave-one-out (LOO) setting.** **a**, Locityper LOO accuracy at 40 Illumina datasets. **b**, Locityper LOO accuracy at 40 simulated short-read datasets. *[Continued on the next page]*

**c**

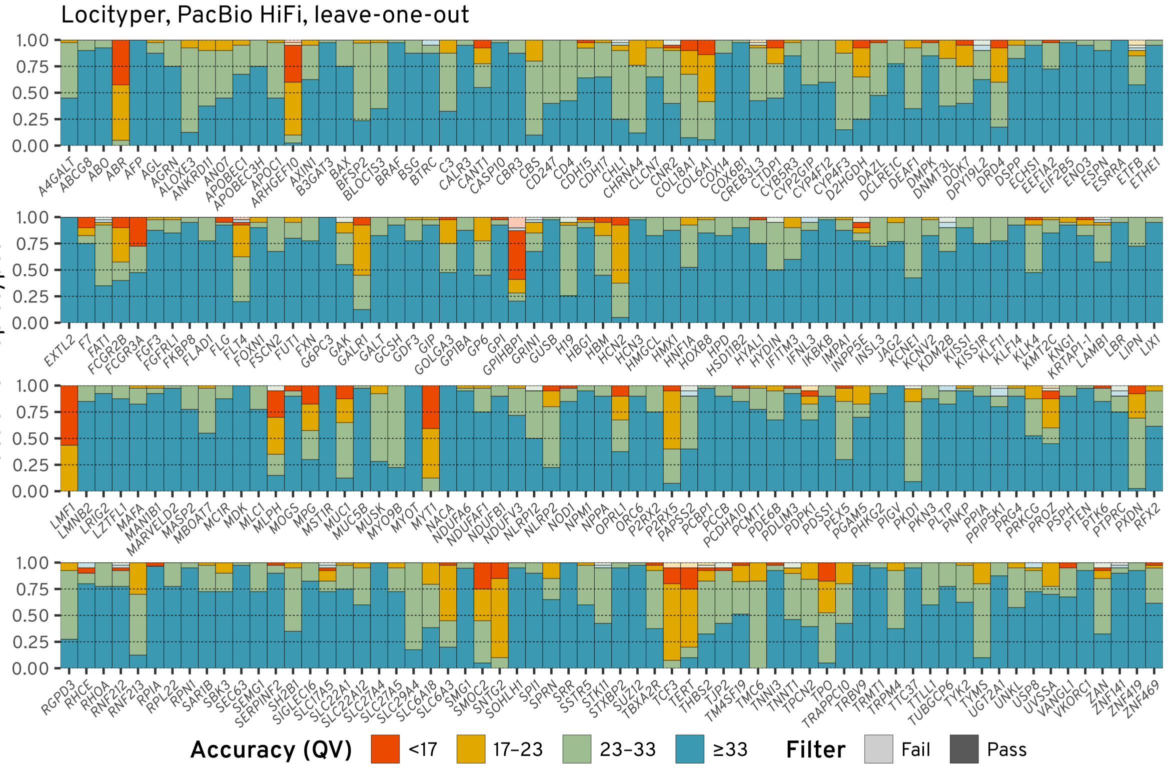

**d**

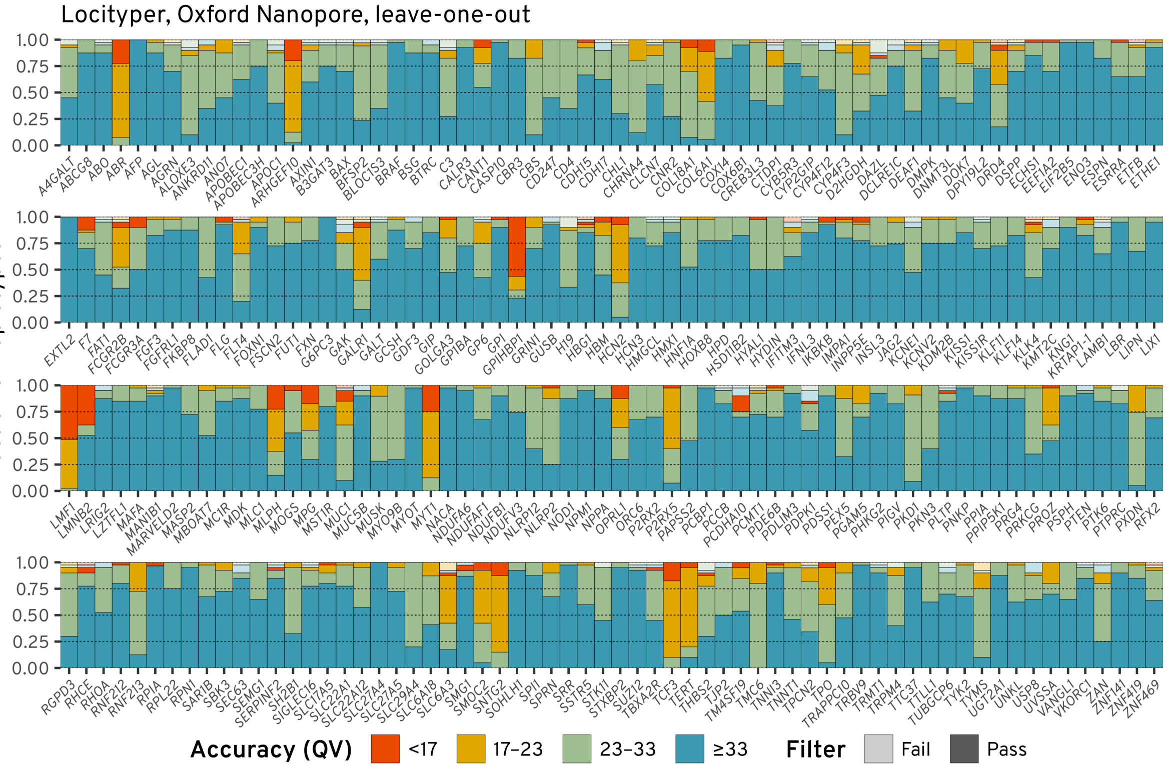

**Supplementary Figure 2.** *(continued)* **c**, Locityper LOO accuracy at 20 PacBio HiFi datasets. **d**, Locityper LOO accuracy at 20 Oxford Nanopore datasets.

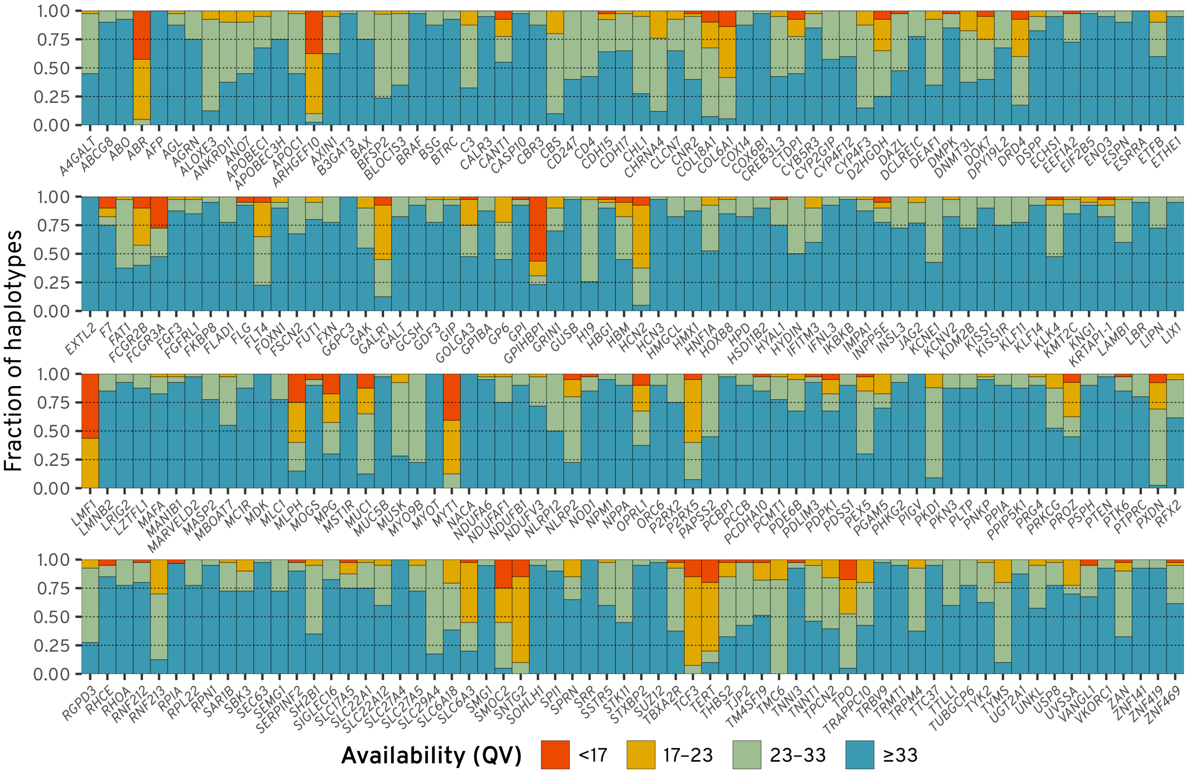

**Supplementary Figure 3. Haplotype availability in the leave-one-out setting.** In the leave-one-out setting, two actual sample haplotypes are removed from the database. This figure shows Phred-scaled divergence (QV) between the actual haplotypes and the closest remaining haplotypes.

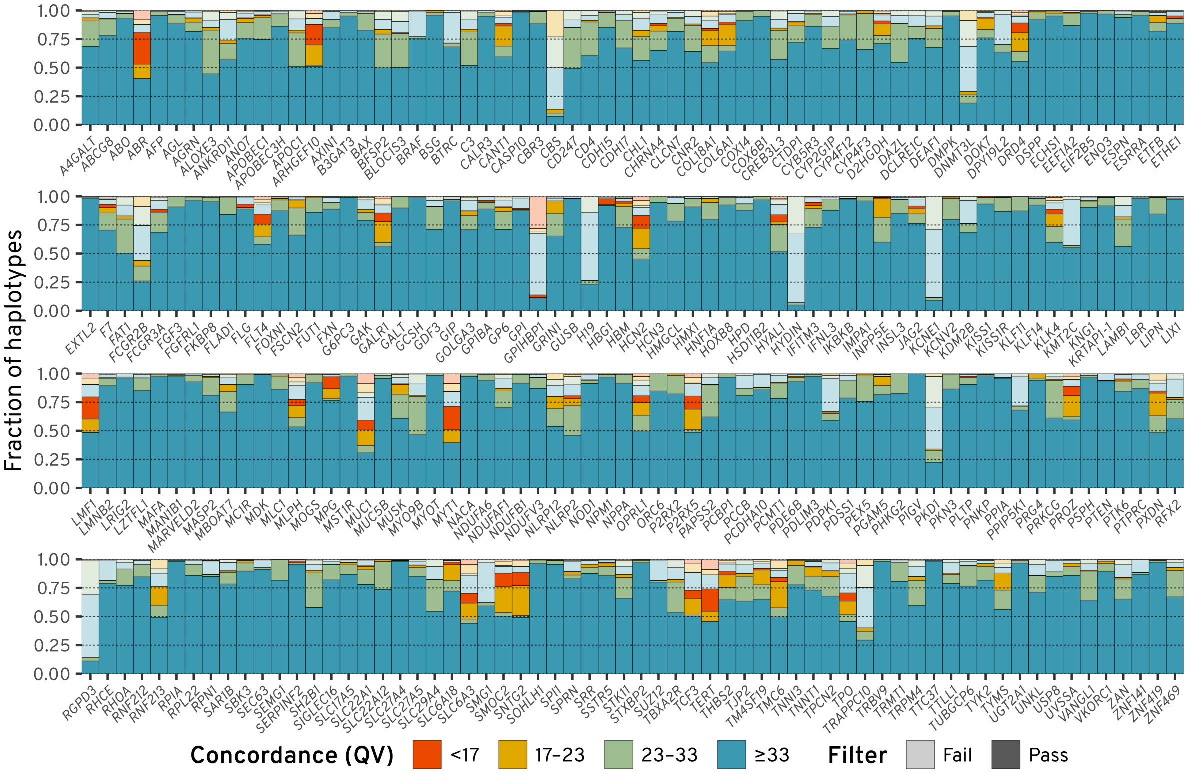

**Supplementary Figure 4. Locityper prediction concordance for 602 trios across 256 challenging medically relevant loci.** Here, trios are considered to fail filtering (semi-transparent colors) if any of the three individual genotypes failed filtering.

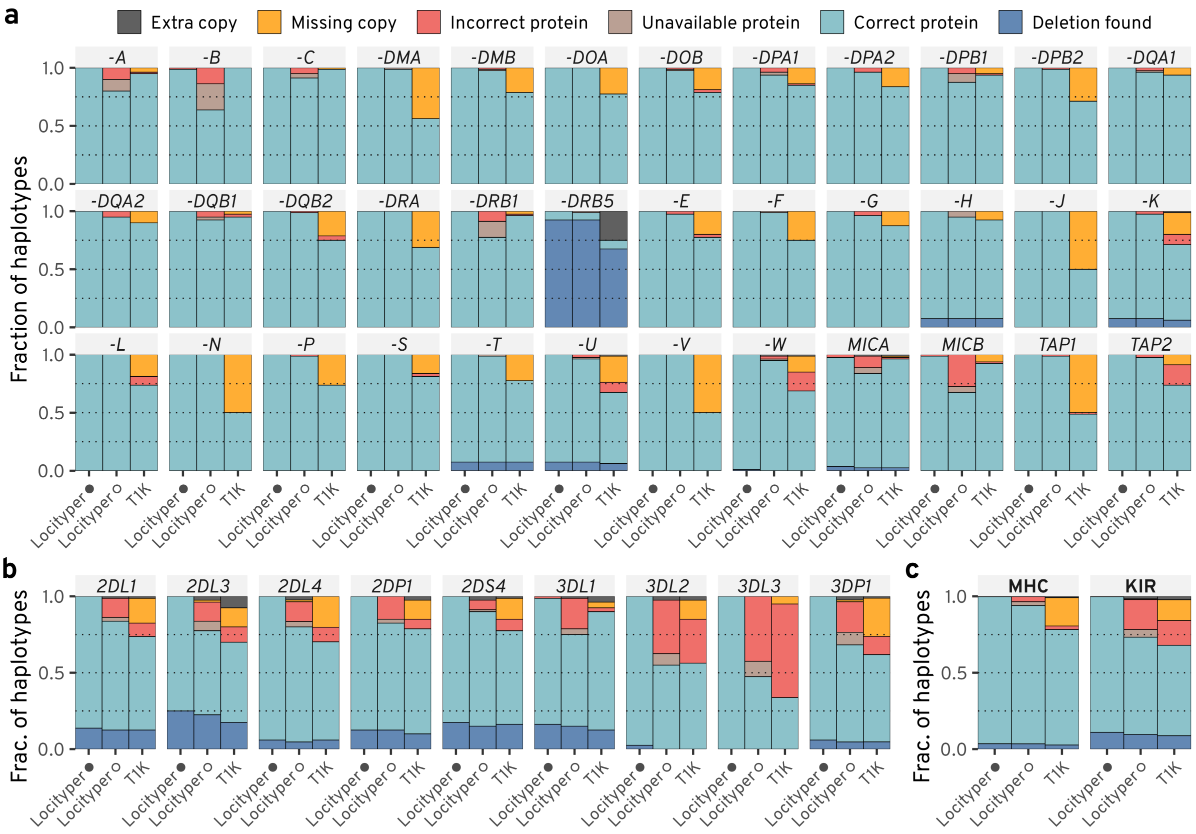

**Supplementary Figure 5. Stratification of HLA/KIR calls.** On each subplot, accuracy is shown for Locityper with the full database (denoted by black circle); Locityper in the leave-one-out (LOO) setting (white circle); and T1K. Genotyping is performed for 40 HPRC samples across 36 (pseudo)genes from the MHC locus (**a**) and 9 (pseudo)genes from the KIR locus (**b**). Panel **c** shows aggregate counts across all genes/pseudogenes from the MHC and KIR loci. T1K/Locityper allele predictions are placed into six categories: *extra copy* for cases when a genotyper called more gene copies than actually present in the locus; *missing copy* when a genotyper failed to call a present gene copy; *(in)correct protein* for predictions where a protein product (second field in the HLA/KIR nomenclature) was called (in)correctly; *unavailable protein* for such Locityper LOO predictions, where true protein product is unavailable in the LOO database and therefore cannot be correctly identified; and *deletion found* for cases when a genotyper correctly identified a missing gene copy.

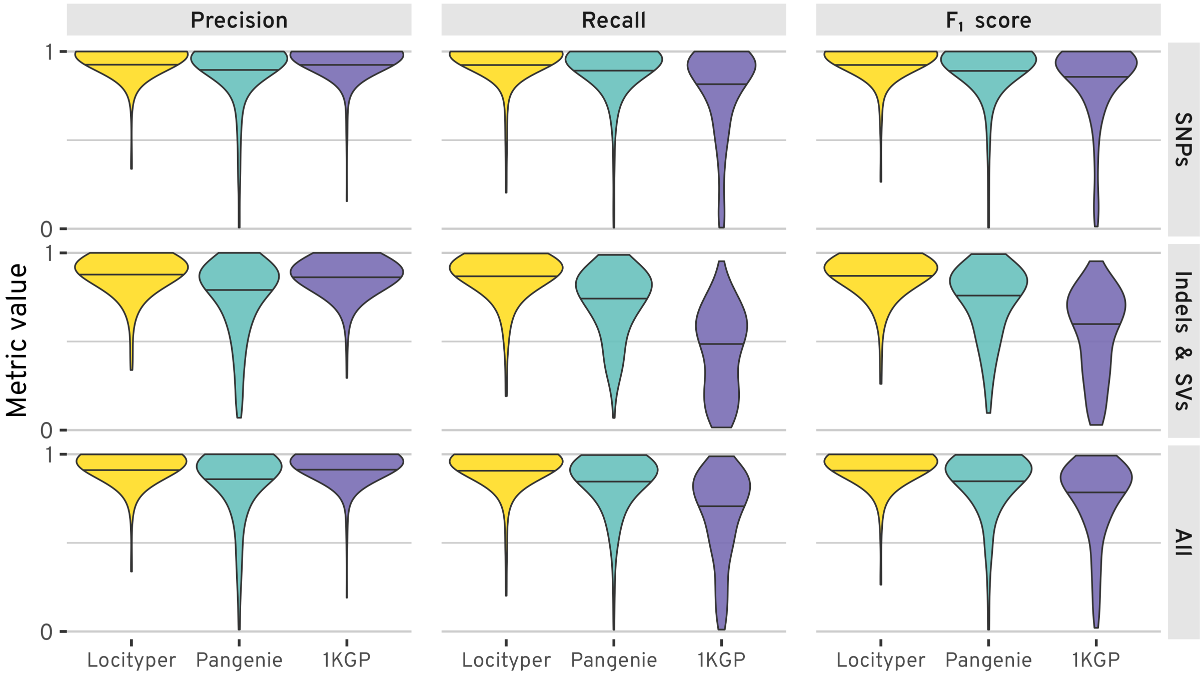

**Supplementary Figure 6. Variant calling accuracy comparison for Locityper, Pangenie, and the 1KGP variant call sets across 256 medically relevant loci.** The figure shows variant calling precision, recall and $F_1$ scores, stratified by the variant type (SNPs; indels and SVs; and all together). Black horizontal lines show median values across all loci. Analyised samples overlap input haplotype databases for both Locityper and Pangenie.

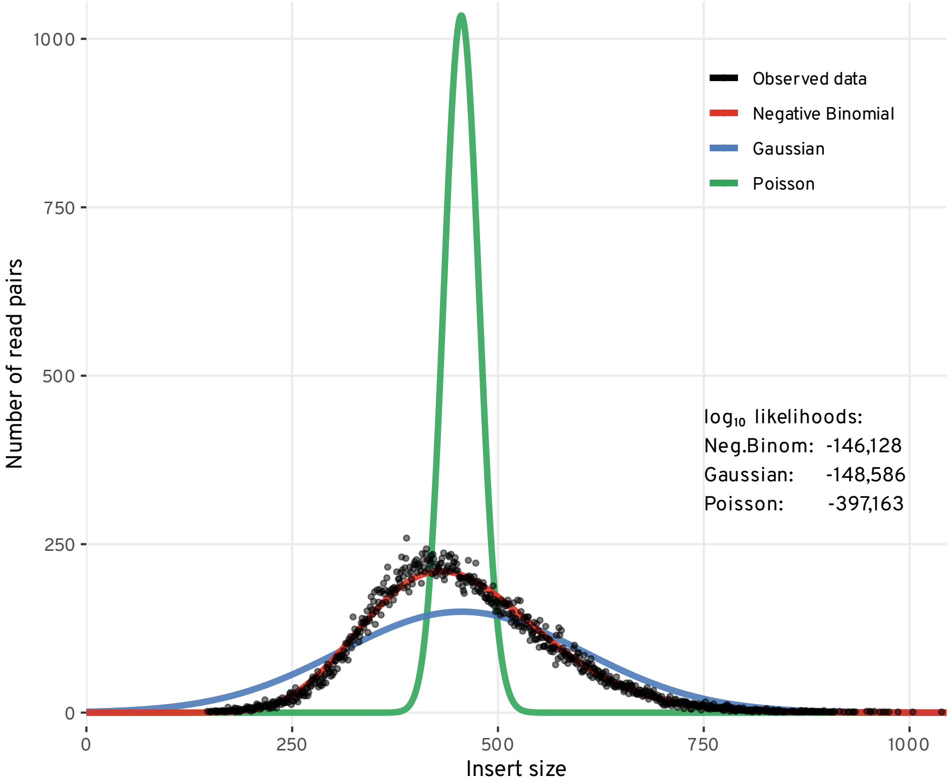

**Supplementary Figure 7. Insert size distribution.** Black dots show observed insert sizes for 55 thousands read pairs from the HG00621 Illumina WGS dataset. Colored lines show three fitted distributions: Negative Binomial (red), Gaussian (blue) and Poisson (green). Fit $\log_{10}$ likelihoods for all distributions are shown on the right of the figure.

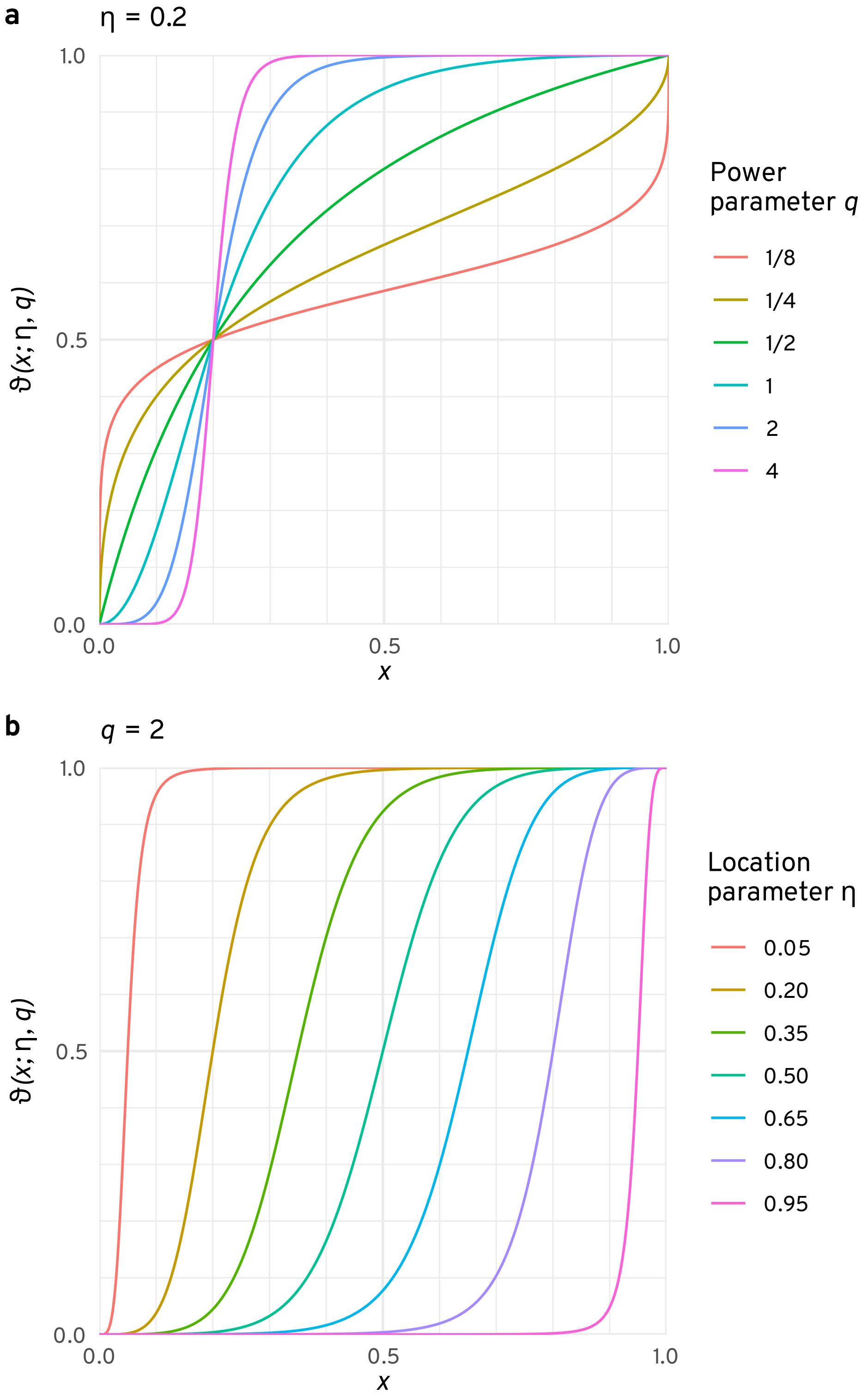

**Supplementary Figure 8. Two-parametric weight function** $\vartheta(x;\eta,q)$ with fixed $\eta = 0.2$ and variable $q$ (**a**) as well as fixed $q = 2$ and variable $\eta$ (**b**).

### 2 Supplementary Tables

**Supplementary Table 1. Locityper target regions.** Description of 312 target loci: 256 challenging medically relevant loci, 25 loci from the MHC cluster, 1 KIR locus, and additional 30 loci covering MUC, CFH and CYP2 loci. The table contains locus coordinates and lengths, as well as all covered protein coding genes and non-protein coding genes or pseudogenes. Genes that overlap a locus, but are not contained in it, are marked with an asterisk.

### 3 Supplementary Methods

#### 3.1 Estimating Negative Binomial parameters

Negative Binomial (NB) parameters $n$ and $\psi$ can easily be calculated based on the sample mean $m$ and variance $v$ using the method of moments:

$$n = \frac{m^2}{v - m}, \quad \psi = \frac{m}{v}. \tag{1}$$

However, in cases when the underlying distribution is similar to the Poisson distribution, observed variance $v$ can become very similar to mean $m$ (or even smaller than $m$). In such cases, parameter $n$ can become very large, or even negative (forbidden under NB definition). For these reasons, we employ $L_1$ regularization on $n$.

Additionally, it has been shown that under Binomial subsampling with rate $s$ (also known as $s$-thinning), distribution $\mathrm{NB}(n, \psi)$ is transformed[1,2] into distribution $\mathrm{NB}\left(n, \frac{\psi}{s+\psi-s\psi}\right)$. Accordingly, we correct read depth distributions according to the subsampling rate $s$, used during WGS dataset preprocessing. To summarize, we estimate NB parameters $n$ and $\psi$ based on sample mean $m$, variance $v$ and subsampling rate $s$:

$$\operatorname*{argmin}_{\substack{n>0 \\ \psi\in(0,1)}} \left(\frac{ns \cdot (1-\psi)}{\psi} - m\right)^2 + \left(\frac{ns \cdot (1-\psi) \cdot (\psi + s - \psi s)}{\psi^2} - v\right)^2 + \lambda n, \tag{2}$$

where $\lambda$ is the regularization parameter ($10^{-5}$ by default).

##### 3.2 Alternative Integer Linear Programming Formulation

Theoretically, locus genotyping problem can be stated in a single ILP statement. For a given ploidy $\pi$ let us define an integer variable $q_a \in \{0, \ldots, \pi\}$ for each haplotype $a \in A$. Then, we can generalize the problem statement from the main text in the following way:

$$\begin{aligned}
\text{Maximize} \quad & \Omega_R \sum_{\mathbf{r} \in R} \sum_{a \in A} \sum_{\mathbf{w} \in L_{\mathbf{r}}^{(a)}} x_{\mathbf{rw}} \cdot \log \mathcal{P}_{\mathbf{rw}} + \Omega_D \sum_{a \in A} \sum_{w \in W^{(a)}} \sum_{d=0}^{D_{\max}} y_{wd} \cdot \zeta_w \cdot \varphi_w(d) \\
\text{Subject to} \quad & \sum_{a \in A} q_a = \pi, \\
& \sum_{a \in A} \sum_{\mathbf{w} \in L_{\mathbf{r}}^{(a)}} x_{\mathbf{rw}} = 1 \qquad \forall \mathbf{r} \in R, \\
& \sum_{d=0}^{D_{\max}} y_{wd} = q_a \qquad \forall a \in A,\ \forall w \in W^{(a)}, \\
& \sum_{\mathbf{r} \in R} \sum_{u \in W^{(\mathbf{g})}} \left( x_{\mathbf{r},wu} + x_{\mathbf{r},uw} \right) - \sum_{d=0}^{D_{\max}} d \cdot y_{wd} = 0 \qquad \forall a \in A,\ \forall w \in W^{(a)}, \\
\text{and} \quad & x_\diamond \in \{0, 1\}, \\
& y_\diamond \in \{0, \ldots, \pi\}, \\
& q_\diamond \in \{0, \ldots, \pi\}.
\end{aligned} \tag{3}$$

Even though this formulation is very flexible and allows for higher ploidy, the large number of variables and complex interactions between them makes the problem almost infeasible for state-of-the-art ILP solvers. Both Gurobi[3] and HiGHS[4] required significantly more time to solve the problem and reached worse likelihoods, compared to the sum time required to solve the ILP problem for each of the locus genotypes. Additionally, generalized problem statement does not allow to directly estimate genotype likelihood for non-primary genotype predictions.
